# TonB dependent uptake of β-lactam antibiotics in the opportunistic human pathogen *Stenotrophomonas maltophilia*

**DOI:** 10.1101/615369

**Authors:** Karina Calvopiña, Punyawee Dulyayangkul, Kate J. Heesom, Matthew B. Avison

## Abstract

The β-lactam antibiotic ceftazidime is one of only a handful of drugs with proven clinical efficacy against the opportunistic human pathogen *Stenotrophomonas maltophilia*, Here, we show that mutations in the energy transducer TonB, encoded by *smlt0009* in *S. maltophilia*, confer ceftazidime resistance. This breaks the dogma that β-lactams enter Gram-negative bacteria only by passive diffusion through outer membrane porins. By confirming cross-resistance of Smlt0009 mutants with a siderophore-conjugated lactivicin antibiotic, we reveal that attempts to improve penetration of antimicrobials into Gram negative bacteria by conjugating them with TonB substrates is likely to select β-lactam resistance in *S. maltophilia*, increasing its clinical threat. Furthermore, we show that *S. maltophilia* clinical isolates that have an Smlt0009 mutation already exist. Remarkably, therefore, β-lactam use is already eroding the potential utility of currently experimental siderophore-conjugated antimicrobials against this species.

## Introduction

*Stenotrophomonas maltophilia* is an important opportunistic human pathogen and clinical isolates are resistant to almost all β-lactam antibiotics because of the production of two β-lactamases: L1, a subclass B3 metallo-β-lactamase and L2, a class A Extended Spectrum β-Lactamase (Gould *et al*., 2006). Production of L1 and L2 is co-ordinately controlled by AmpR, a LysR-type transcriptional activator and induced during β-lactam challenge of cells (Okazaki & Avison, 2008). Despite this, many *S. maltophilia* clinical isolates remain susceptible to the β-lactam ceftazidime because it is a relatively poor substrate for these two enzymes (Calvopina *et al*., 2017). However, mutants that have acquired ceftazidime resistance can easily be identified in the laboratory, and ceftazidime resistant isolates are commonly encountered in the clinic. In many cases, these mutants hyperproduce L1 and L2 (Okazaki & Avison, 2008, Talfan *et al*., 2013, Calvopina & Avison, 2018) but we have previously identified ceftazidime resistant mutants that did not hyperproduce β-lactamase (Gould & Avison, 2006). It was hypothesised that these mutants might have reduced accumulation of ceftazidime (Talfan *et al*., 2013). The primary non-β-lactamase mediated mechanisms of β-lactam resistance in similar non-fermenting bacteria such as *Pseudomonas aeruginosa* are increased efflux and reduced outer membrane permeability due to a reduction in the production of outer membrane porins (Castanheira *et al*., 2014). In Gram negative bacteria generally, tripartite outer membrane porins are considered the only site of entry for β-lactams, and reduced porin levels can reduced β-lactam susceptibility in many species (Pfeifer *et al*., 2010). No other way of entry has previously been suggested unless the β-lactam is conjugated to a catechol siderophore, in which case a TonB-dependent uptake system is used (Livermore, 1987). The aim of the work reported below was to identify the mechanism of non-β-lactamase mediated ceftazidime resistance in *S. maltophilia*. In so doing, we have broken the dogma that states that β-lactams can only enter Gram-negative bacteria through trimeric outer membrane porins via passive diffusion.

## Results and Discussion

Around 50% of ceftazidime resistant mutants selected from *S. maltophilia* clinical isolate K279a do not hyperproduce β-lactamase (Talfan *et al*., 2013). To identify the mechanism involved, we selected additional ceftazidime resistant mutants from K279a. Mutants expressing basal β lactamase activity (i.e. in the absence of β-lactam antibiotic) similar to K279a (∼0.02 nmol of nitrocefin hydrolysed.min^-1^.µg^-1^ of extracted protein) were taken forward for study. Of these, mutants M1 and M52 are exemplars. They are not β-lactamase hyperproducers (**Table 1**) but β lactam susceptibility was reduced, as shown by an observed reduction in the inhibition zone diameter around various β-lactam discs. However, where non-β-lactams were tested, the impact on susceptibility of the mutations was minimal (**Figure 1**).

**Table 1.** Comparison of MICs (mg.L^-1^) of ceftazidime and lactivicin derivatives against *S. maltophilia* ceftazidime and lactivicin mutants and the levels of β-lactamase produced.

| Strain | Mean β-lactamase activity ±SEM | MIC of Ceftazidime | MIC of LTV-13 | MIC of LTV-17 |
| --- | --- | --- | --- | --- |
| K279a | 0.02±0.004 | 4 | 64 | 0.03 |
| M1 | 0.02±0.002 | 256 | 128 | 0.5 |
| M52 | 0.04±0.013 | 256 | 128 | 0.5 |
| KLTV | 0.05±0.005 | 256 | 64 | 0.25 |
| K279a Δsmlt0009 | 0.01±0.002 | 128 | 128 | 0.5 |

β-Lactamase activity was determined using nitrocefin hydrolysis (nmol.min<sup>-1</sup>. μg<sup>-1</sup>) in cell extracts from bacteria grown in the absence of antibiotic.
Shaded values represent a more than two doubling reduced susceptibility in reference to K279a and in the case of ceftazidime, shading show clinical resistance according to CLSI breakpoints.

**Figure 1.**
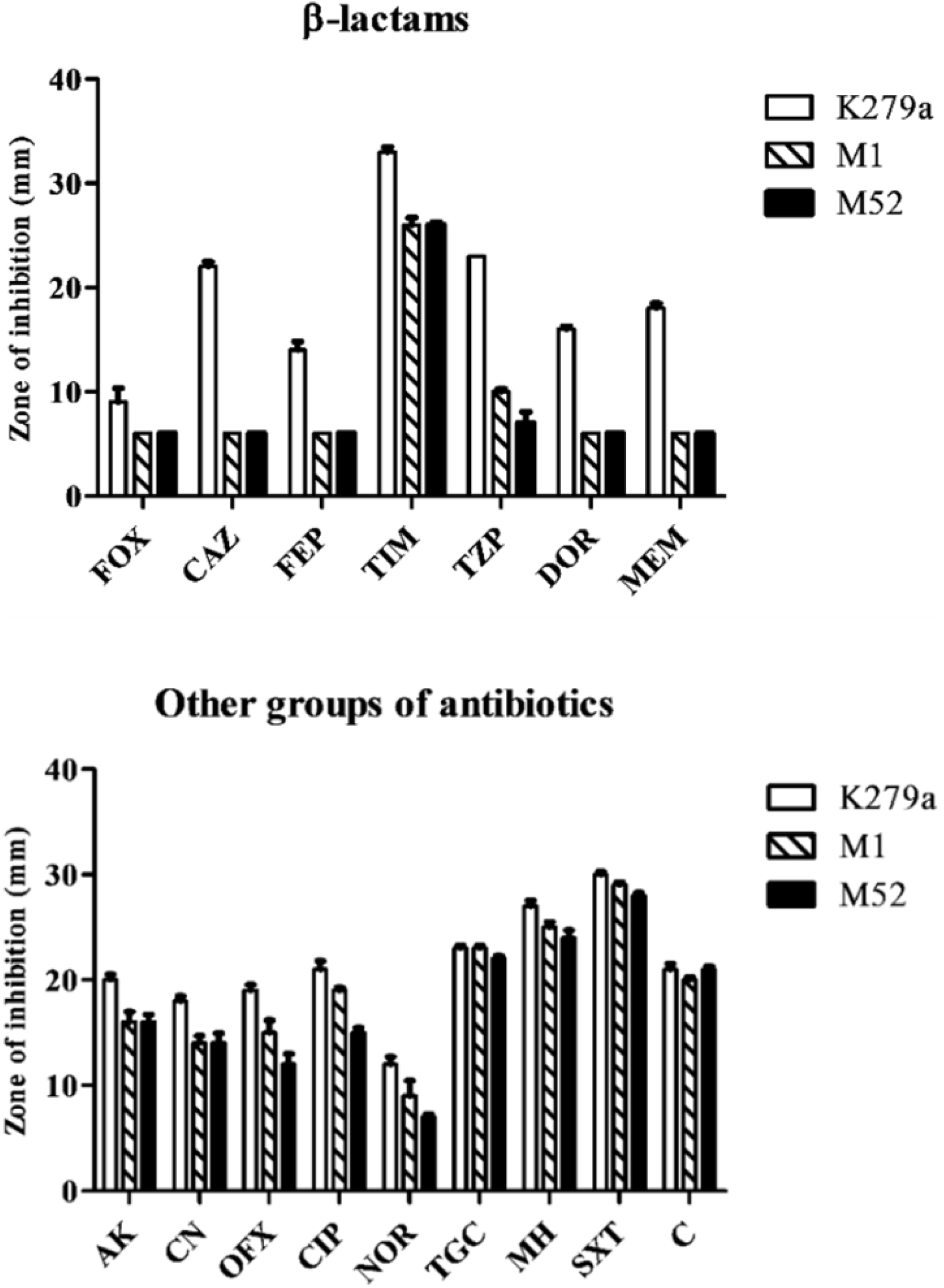
Antibiotic susceptibilities of ceftazidime resistant mutants versus K279a. Growth inhibition zone diameters (mm) of ceftazidime resistant mutants (M1 and M52) in comparison with the parental strain (K279a). Smaller zone diameters mean reduced susceptibility. The following antibiotics were tested: (**A**) β-lactams; cefoxitin (FOX 30 µg), ceftazidime (CAZ 30 µg), cefepime (FEP 30 µg), ticarcillin-clavulanate (TIM 85 µg), piperacillin-tazobactam (TZP 110 µg), doripenem (DOR 10 µg), meropenem (MEM 10 µg). (B) non-β-lactams; amikacin (AK 30 µg), gentamicin (CN 10 µg), ofloxacin (OFX 5 µg), ciprofloxacin (CIP 5 µg), norfloxacin (NOR 10 µg), tigecycline (TGC 15 µg), minocycline (MH 30 µg), trimethoprim-sulfamethoxazole (SXT 25 µg), chloramphenicol (C 30 µg). Zones of inhibition are reported as mean values, n=3. Error bars represent standard error of the mean (SEM). Zone diameters are measured across the disc, so the minimum zone diameter is 6 mm, which is the diameter of the disc.

Whole genome sequencing was performed to identify the mutations present in ceftazidime resistant mutants M1 and M52. Only one gene was found to be mutated in each. It was the same in both: *smlt0009*, annotated in the K279a genome sequence as encoding a ‘putative proline-rich TonB energy transducer protein’ (Crossman *et al*., 2008). The mutation was confirmed using high fidelity PCR sequencing. In both M1 and M52, a proline rich region in Smlt0009 situated at around 70 amino acids into the 222 amino acid protein was shortened but there was no frameshift (**Figure 2**). Assuming this shortening impairs protein function, and to confirm the role of this impairment in ceftazidime resistance, *smlt0009* was insertionally inactivated in K279a using a suicide gene replacement methodology. K279a Δ*smlt0009* was confirmed to be ceftazidime resistant (**Table 1**).

**Figure 2.**
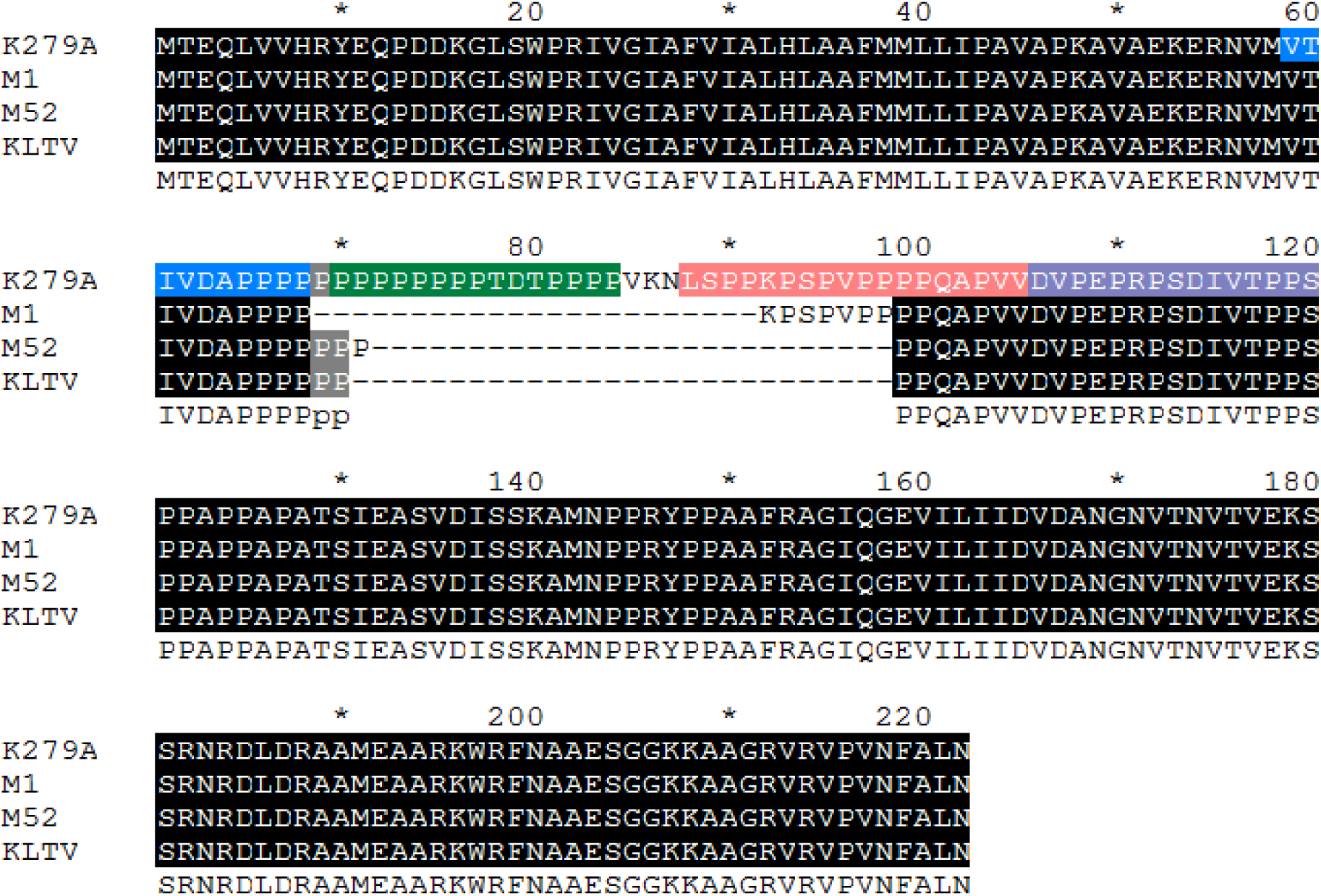
Sequence alignment of Smlt0009 putative proline-rich TonB energy transducer protein in ceftazidime resistant mutants versus K279a. Alignment of translated high fidelity PCR sequences that confirmed mutation in the proline-rich region in M1, M52 and KLTV. Key residues present in *E.coli* TonB (Kohler *et al.*, 2010) as compared with Smlt0009 that dictate periplasmic spanning distance are highlighted: blue bar corresponds to 2.9nm, the green bar to 4nm, pink bar 4.6nm and purple bar 3.3nm. Alignment was performed with CLUSTAL Omega and GeneDoc.

To understand more about the phenotype of M1 and M52, whole envelope proteomics was performed in comparison with K279a. This confirmed that the β lactamases L1 and L2 are not overproduced. However, 162 proteins were identified that are significantly up or down regulated in both M1 and M52 relative to K279a; 83 are downregulated in both and 79 upregulated in both (**Table S1**). Within the group of downregulated proteins, Smlt0009 (Uniprot: B2FT87) itself was 1.8 fold downregulated in M1, and 2.5-fold in M52 when compared with K279a (**Figure 3A**). Shortening of the Smlt0009 proline rich region in M1 and M52 is not expected to block production of the protein, but the mutated, presumably less active, protein may be less stable than wild-type explaining this downregulation. Proteomics for K279a Δ*smlt0009* (**Table S2**) confirmed total loss of Smlt0009 in this case (**Figure 3A**). Amongst proteins upregulated in M1, M52 (and in K279a Δ*smlt0009*) were proteins with the Uniprot accession numbers B2FHQ4, encoded by *entB, smlt2820* (**Figure 3B**), B2FRE6 (*fepC, smlt2356*) and B2FRE7 (*fepD, smlt2357*) (**Tables S1; S2**). These upregulated Fep proteins are involved in siderophore production in *S. maltophilia* (Nas & Cianciotto, 2017) and siderophore production was found to be increased in M1 and M52 and K279a Δ*smlt0009* relative to K279a, as predicted from the proteomics (**Figure 3C**).

**Figure 3.**
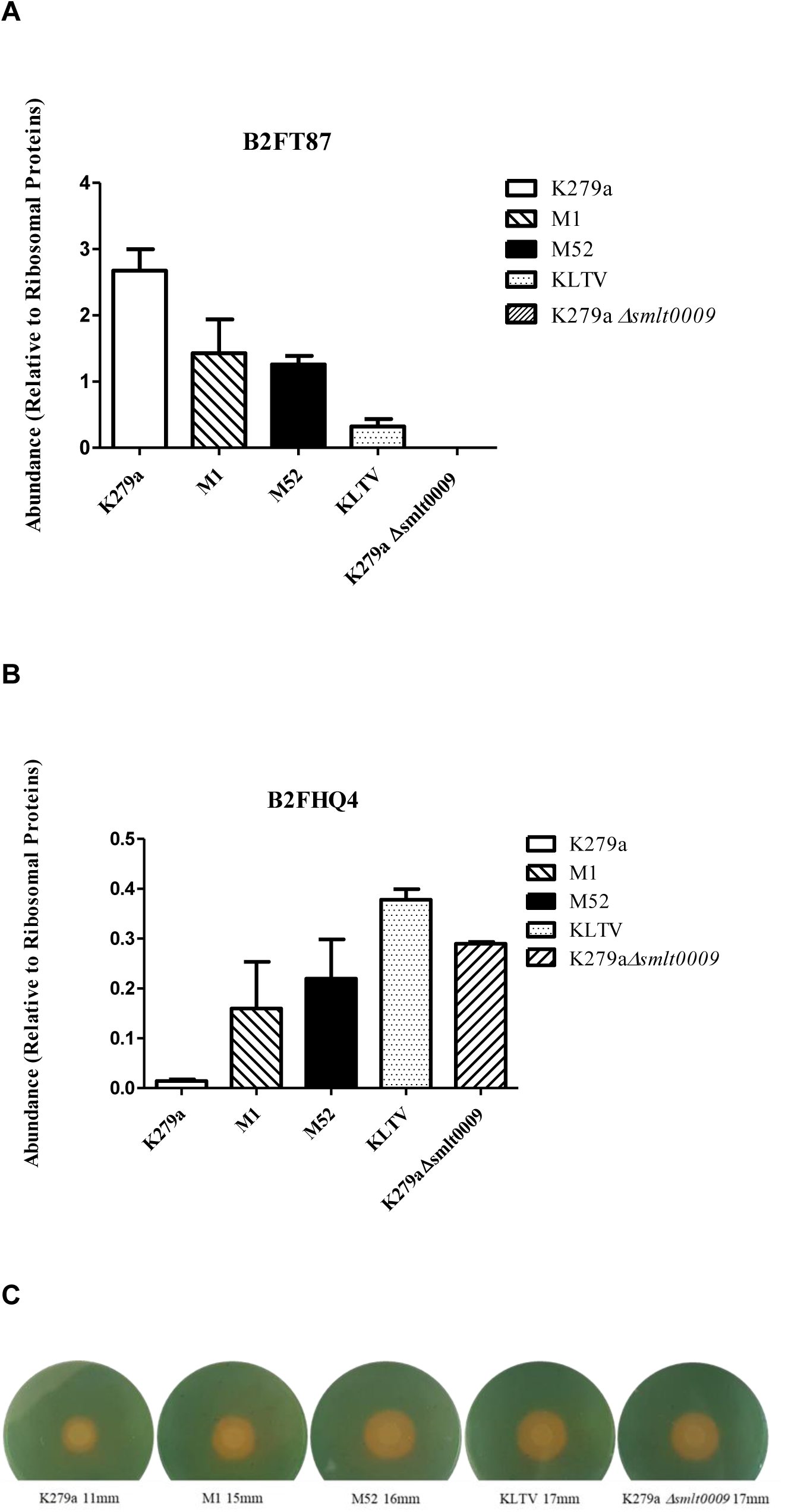
Abundance of key proteins and siderophore production in ceftazidime resistant mutants versus K279a. Protein abundance data for (**A**) the TonB energy transducer protein Smlt0009 (Uniprot: B2FT87) and (**B**) EntB (Uniprot: B2FH84) were each normalised using the average abundance of 30S and 50S ribosomal proteins in each sample. Values are reported as mean +/-Standard Error of the Mean (*n*=3). In each case the change relative to K279a in each mutant is statistically significant (p<0.05). Full Proteomics data are shown **Tables S1, S2 and S3**. (**C**) Siderophore production assay. Diameter values show diffusion of siderophore after spotting 10 µL of a PBS washed bacterial suspension (OD_600_ 0.2) onto a modified CAS agar. Values are reported as mean of three biological repeats; the images are representative of these three experiments.

Once iron-scavenging siderophores are exported by a bacterium, iron-siderophore-complex import requires a TonB complex formed by a proline rich TonB energy transducer protein with ExbB and ExbD, which interacts with one or more ligand-gated porins (LGPs). Specificity occurs because TonB only interacts with LGPs that have bound substrate (Wilson *et al*., 2016, Klebba, 2016). In this way, proton motive force, generated in the inner membrane, is transduced by ExbBD – Smlt0010 and Smlt0011 in *S. maltophilia* (Crossman *et al*., 2008) – to cause rotational motion of the N-terminus of the TonB energy transducer (Smlt0009) and specific opening of any LGP that has bound ligand, ultimately driving ligand import (Klebba, 2016).

Ceftazidime resistant mutants M1 and M52 have mutations in this proline rich TonB energy transducer protein, Smlt0009, so TonB complex dependent import of all LGP ligands is likely to be reduced. These mutants also have upregulation of proteins involved in siderophore production (**Table S1; Figure 3B**), leading to observed enhanced siderophore production (**Figure 3C**). One hypothesis to explain this is that loss of Smlt0009 activity impedes iron siderophore-complex import, which increases siderophore production as a response to the resulting iron starvation.

In some bacteria, TonB complexes participate in the import of LGP-dependent ligands in addition to iron-siderophore complexes. In fact, in the environmental species *Xanthomonas campestris*, only 15% of LGPs are involved in iron-siderophore-complex import (Schauer *et al*., 2008). *S. maltophilia* Smlt0009 shares 50% identity with the TonB energy transducer protein from the closely related species *X. campestris*. Interestingly, of 162 proteins differently regulated in M1 and M52, 19 are putative TonB-dependent LGP proteins (**Table S1**). Apparently, M1 and M52 are responding to a breakdown in TonB-dependent energy transduction associated with the import of many diverse ligands.

In terms of ceftazidime resistance in M1 and M52, we hypothesise that in *S. maltophilia*, β lactams are TonB dependent substrates. Thus, mutations in the proline rich region of Smlt0009 reduce energy dependent-ceftazidime import. This is the first time that β lactam entry via a TonB dependent mechanism has been proposed in any bacterium. However, it is interesting to note that, unlike all other pathogens studied previously, outer membrane passive diffusion porin loss has never been seen to be involved in β-lactam resistance in *S. maltophilia* (Sanchez, 2015) which supports the existence of a novel import mechanism in this species.

To test our hypothesis that reduction of Smlt0009 activity reduces ceftazidime import in *S. maltophilia*, we tested envelope permeability to a DNA-intercalating Hoescht dye in the presence of ceftazidime. In K279a, permeability to the dye (and so the rate of increase in cellular fluorescence following binding of the dye to DNA) reduced in the presence of ceftazidime, which means both antibiotic and dye are competing for the same general uptake system(s) (**Figure 4A**). This reduction in permeability is not caused by cell death because the concentration of ceftazidime chosen does not significantly impact on cell growth during mid-exponential phase where cells are harvested to test permeability (**Figure 4B**). Smlt0009 mutant M1, chosen as an exemplar, is less permeable to the dye than K279a, presumably because entry of the dye is at least in part TonB dependent, but most importantly, in M1, ceftazidime no longer competes with the dye for entry to the cell (**Figure 4B**). Therefore, we conclude that Smlt0009 mutation reduces ceftazidime uptake, and that this is the mechanism of ceftazidime resistance.

**Figure 4.**
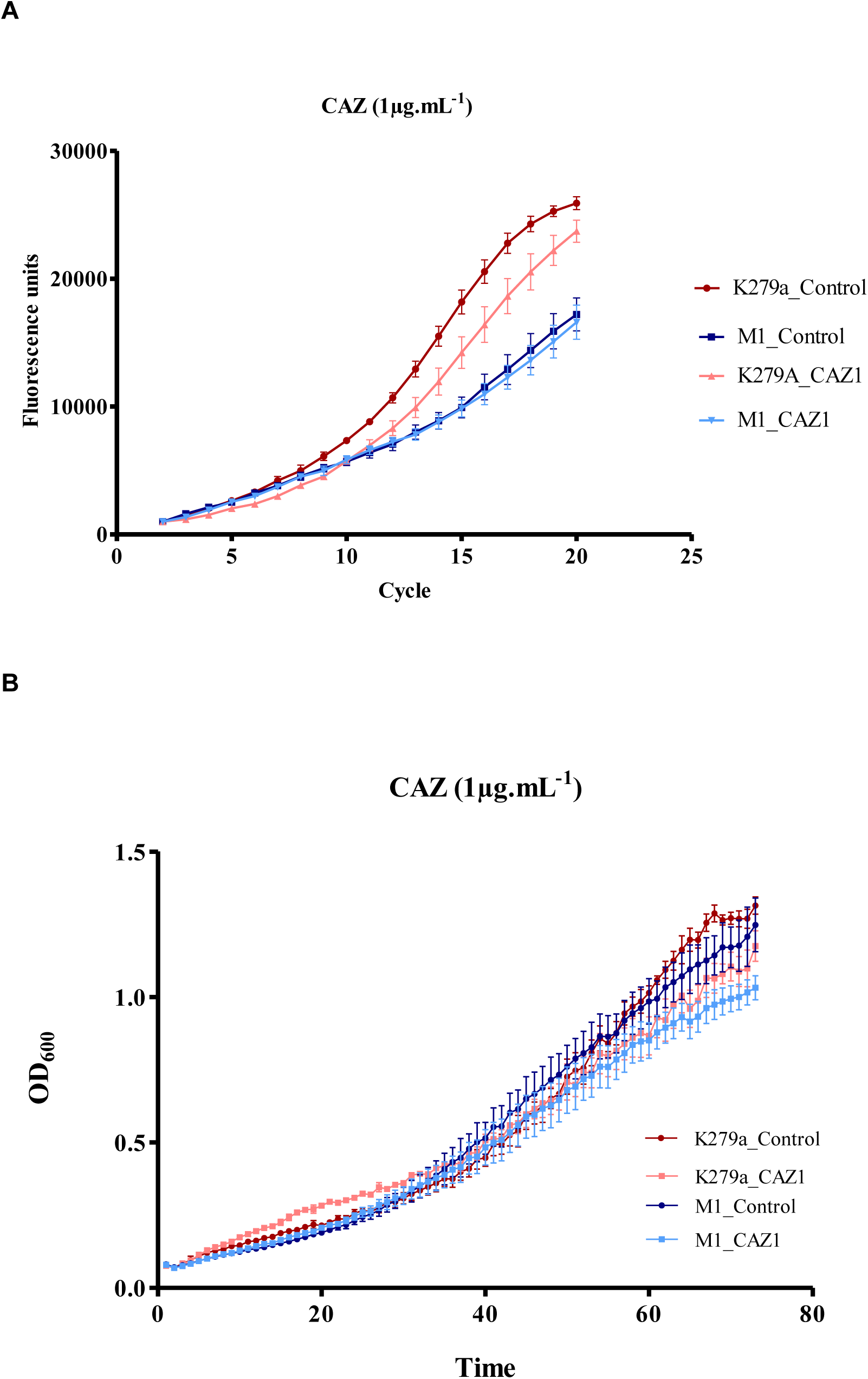
Fluorescent dye accumulation over time in ceftazidime resistant mutant M1 versus K279a in the presence and absence of ceftazidime. (**A**) Rate of fluorescent dye accumulation in K279a (control-red) is reduced in the presence of 1 µg.mL^-1^ ceftazidime (CAZ1 - pink). Rate of fluorescent dye accumulation in M1 in the absence of ceftazidime (control - dark blue) is lower than in K279a, but this is not reduced by the presence of 1 µg.mL^-1^ ceftazidime (CAZ1 - light blue). (**B**) Growth curves in NB in the absence of ceftazidime (K279a, control - red and M1, control - dark blue) are not dramatically altered in the presence of ceftazidime at 1 µg.mL^-1^ (K279, CAZ1 - pink and M1 CAZ1 - light blue) over a 73 cycle (12h) incubation period. Each curve plots mean data for three biological replicates with, in **B**, four technical replicates for each biological replicate. Error bars represent standard error of the mean.

Siderophore-conjugation has been used as a way of increasing the penetration of cephalosporins and monobactams into Gram-negative bacteria by hijacking the TonB dependent uptake system (Kline *et al*., 2000, Choi & McCarthy, 2018). Indeed, recently we have shown that siderophore conjugation of the γ-lactam antibiotic lactivicin (to create LTV-17) dramatically improves potency against *S. maltophilia* (Calvopina *et al*., 2016). As expected given TonB dependence of LTV-17 uptake (Starr *et al*., 2014), ceftazidime resistant Smlt0009 mutants M1 and M52 also have reduced susceptibility to LTV-17, as does K279a Δ*smlt0009* where in each case the MIC increased to ≥0.25 µg.mL^-1^ (**Table 1**). A single-step mutant (KLTV) with reduced susceptibility to LTV-17 was next selected from K279a and we found that the mutant is also resistant to ceftazidime (**Table 1**). KLTV whole envelope proteomics showed very similar changes to those observed in M1 and M52 (**Table S3**). Specifically, in KLTV, like M1 and M52, there is downregulation of Smlt0009 and upregulation of the EntB siderophore biosynthesis enzyme and siderophore overproduction (**Figure 3**). WGS confirmed shortening of the proline-rich region in Smlt0009 in KLTV (**Figure 2**). TonB mutations are known to increase susceptibility to siderophore conjugated antimicrobials but have never previously been reported to affect β-lactam susceptibility (Hassett *et al*., 1996, Tomaras *et al*., 2013, Moynie *et al*., 2017). Importantly, *S. maltophilia* Smlt0009 mutants do not have reduced susceptibility to the non-siderophore conjugated parent lactivicin, LTV-13 (**Table 1**), even though this γ-lactam is structurally related to the β-lactams (Starr *et al*., 2014). This shows that TonB-dependent uptake of β-lactams is specific and there is minimal affinity for γ-lactams.

Finally, we turned to our world-wide collection of 22-phylogenetic group A *S. maltophilia* clinical isolates (Gould *et al*., 2006) against which we measured the MICs of LTV-17 and LTV-13 (**Table 2**). One isolate, number 31, stood out as having reduced susceptibility to LTV-17 (MIC = 0.25 µg.mL^-1^) without altered susceptibility to LTV-13, a phenotype shared with K279a Smlt0009 mutants (**Tables 1, 2**). Of the tested clinical isolates had the same predicted sequence for Smlt0009 as K279a, based on PCR sequencing; eight isolates had N169S plus A209T variants of this sequence, but given it is so common this is highly likely to be random genetic drift. Isolate number 31, with reduced LTV-17 susceptibility had an insertion of a single proline in the proline-rich region of Smlt0009 (**Table 2**).

**Table 2.** MICs (mg.L^-1^) of lactivicin derivatives against *S. maltophilia* clinical isolates with different Smlt0009 sequences.

| Isolate | Smlt0009 sequence<br>(as compared with K279a) | LTV-13 MIC | LTV-17 MIC |
| --- | --- | --- | --- |
| K279a | Wild-type (by definition) | 64 | 0.03 |
| 10 | N169S, A209T | 128 | 0.03 |
| 12 | Wild-type | 64 | 0.03 |
| 14 | Wild-type | 64 | 0.03 |
| 16 | Wild-type | 64 | 0.03 |
| 17 | Wild-type | 64 | 0.03 |
| 19 | Wild-type | 128 | 0.06 |
| 21 | Wild-type | 64 | 0.03 |
| 22 | Wild-type | 128 | 0.03 |
| 23 | N169S, A209T | 256 | 0.03 |
| 26 | N169S, A209T | 64 | 0.03 |
| 27 | N169S, A209T | 128 | 0.06 |
| 28 | Wild-type | 64 | 0.03 |
| 29 | Wild-type | 64 | 0.03 |
| 30 | N169S, A209T | 128 | 0.03 |
| 31 | Insertion of Proline between<br>P69 and P70 | 128 | 0.25 |
| 32 | N169S, A209T | 128 | 0.13 |
| 35 | N169S, A209T | 64 | 0.06 |
| 36 | Wild-type | 64 | 0.03 |
| 37 | Wild-type | 64 | 0.06 |
| 39 | Wild-type | 128 | 0.03 |
| 40 | N169S, A209T | 64 | 0.03 |
| 43 | Wild-type | 128 | 0.06 |

According to our records, isolate number 31 was from a patient being treated in an intensive care unit in a Brazilian hospital in 2003. It was collected as part of the SENTRY antimicrobial surveillance programme (Toleman *et al*., 2007). Whilst siderophore-conjugated antimicrobials have been in experimental use since the 1980s there is no reason to believe that this isolate has ever been exposed to such a compound. We must conclude, therefore that this mutation has been selected by β-lactam use. Remarkably, however, isolate number 31 carries an *ampD* loss of function mutation and hyper-produces both the L1 and L2 β-lactamases, which is enough to give pan β-lactam resistance without any additional mechanism (Gould *et al*., 2006, Talfan *et al*., 2013, Calvopina *et al*., 2017). It is possible, however, that some combination of β-lactams would still be effective against a β-lactamase hyper-producer, and the use of such a combination might select for this additional β-lactam resistance mechanism. And certainly, isolate number 31 is unusual in its resistance to ceftazidime/β-lactamase inhibitor combinations (Calvopina *et al*., 2017), so combination therapy including a β-lactam/β-lactamase inhibitor might have selected for this mutation even in a background of β-lactamase hyper-producer. Whatever the specifics of selection in this case, we have demonstrated the existence of *S. maltophilia* clinical isolates with mutations in the TonB energy transducer which have reduced susceptibility to β-lactams and siderophore-conjugated antimicrobials. Uncovering this unforeseen cross-resistance phenotype may well suggest a reassessment of the use of β-lactams, alone or in combination with other agents, for the treatment of *S. maltophilia* infections unless there is no alternative for fear of eroding the future potential of siderophore-conjugated antimicrobials as agents to treat infections caused by this species.

## Experimental

### Bacterial isolates and materials

*S. maltophilia* clinical isolates used originated from the SENTRY antimicrobial resistance survey and have been previously described (Toleman *et al*., 2007) plus isolate K279a (Avison *et al*., 2000). All growth media were from Oxoid. Chemicals were from Sigma, unless otherwise stated. LTV-13 was re synthesized according to the literature protocol and kindly provided by Prof. C. Schofield, University of Oxford (Starr *et al*., 2014). LTV-17 was kindly supplied by Pfizer.

### Selection of resistant mutants

K279a ceftazidime resistant mutants were selected after exposure of lawns of bacteria to 30 µg ceftazidime discs on Muller-Hinton Agar (MHA) by picking the colonies within the zone of inhibition after using a bacterial suspension that was 100-fold higher than the recommended value according to the CLSI guidelines (CLSI, 2012). Mutants with reduced susceptibility to LTV-17 were selected by plating 100 µL of an overnight culture grown in Nutrient Broth (NB) on MHA containing increasing concentrations of LTV-17. Colonies from the highest LTV-17 concentration plate where growth was seen were picked.

### Siderophore Detection

100 µL of an overnight culture in Cation-Adjusted Muller-Hinton Broth (CA-MHB) was used to set up a fresh subculture in 10 mL of CA-MHB which was then incubated until the OD_600_ reached 0.5. Cells were centrifuged (4,000 × *g*, 10 min) and the resulting pellet was resuspended in 10 mL of Phosphate Buffered Saline (PBS) and centrifuged again (4,000 × *g*, 10 min). The supernatant was discarded, and the pellet was again resuspended in fresh PBS (10 mL) and centrifuged (4,000 × *g*, 10 min). This washed bacterial pellet was then diluted in PBS to prepare a bacterial suspension of OD_600_ 0.2. Ten microliters of the bacterial suspension were spotted on Chrome Azurol S (CAS) agar. CAS agar was made up mixing up 90 mL of MHA and 10 mL of freshly made CAS solution. 100 mL of the CAS solution was made up based on the following description: 60.5 mg of CAS in 50 mL of water, 72.9 mg of hexadecyltrimethyl ammonium bromide in 40 mL of water, and 10 mL of 1 mM FeCl_3_, 10 mM HCl) (Garcia *et al*., 2012). CAS agar control included 100 µM FeCl3 where no colour change was expected.

### Determining minimal inhibitory concentrations (MICs) of antimicrobials and disc susceptibility testing

The CLSI protocol was followed for disc susceptibility testing (CLSI, 2006). The clearance zone was measured after 20 h of incubation and bacteria reported as susceptible or resistant according to CLSI published breakpoints, where available (CLSI, 2017).

MICs were determined using CLSI broth microtitre assays (CLSI, 2012) and interpreted using published breakpoints (CLSI, 2017). Briefly, a PBS bacterial suspension was prepared to obtain a stock of OD_600_=0.01. The final volume in each well of a 96-well cell culture plate (Corning Costar) was 200 µL and included 20 µL of the bacterial suspension. Bacterial growth was determined after 20 h of incubation by measuring OD_600_ values using a POLARstar Omega spectrophotometer (BMG Labtech).

### β-lactamase assays

100 µL of an overnight NB culture was diluted in 10 mL of NB and incubated at 37°C with shaking until OD600 was 0.4. Cells were pelleted by centrifugation (4,000 × *g*, 10 min) and pellets resuspended in 100 µL of BugBuster (Ambion). Pellets were transferred to 1.5 mL microtube (Eppendorf) before rocking at 70 rpm for 30 min at room temperature. Cell debris and unlysed cells were pelleted by centrifugation (13,000 × *g*, 5 min) and the supernatant retained as a source of crude cell protein. Protein concentrations in cell extracts were determined using the BioRad protein assay dye reagent concentrate according to the manufacturer’s instructions. β-Lactamase activity in crude cell extracts was determined using a POLARstar Omega plate spectrophotometer (BMG Labtech). Nitrocefin (40 μM) solution was used as a substrate, prepared in 0.2 µm syringe-filtered assay buffer (60 mM Na_2_HPO_4_·7H_2_O pH 7.0, 40 mM NaH_2_PO_4_·H_2_O, 10 mM KCl, 1 mM MgSO_4_·7H_2_O, 100 µM ZnCl_2_). Nitrocefin hydrolysis assays were performed in Corning Costar 96-well flat-bottomed cell culture plates with a combination of 1 µL of cell extract and 179 µL of nitrocefin solution. Product accumulation was measured at 482 nm for 5 min or until the end of the linear phase of the reaction. Final β-lactamase activity (nmol.min^-1^.µg^-1^ of protein in cell extract) was calculated via change in absorbance per minute taken from the linear phase of the reaction in Omega Data Analysis. An extinction coefficient of 17400 M^-1^cm^-1^ was used for nitrocefin. The path length for liquid in a well in the 96-well plate was set at 0.56 cm.

### Fluorescent Hoescht (H) 33342 dye accumulation assay

Envelope permeability in living bacteria was tested using a standard dye accumulation assay protocol (Coldham *et al*., 2010) where the dye only fluoresces if it crosses the entire envelope and interacts with DNA. Overnight cultures (in NB) at 37°C were used to prepare NB subcultures, which were incubated at 37°C until a 0.6 OD_600_ was reached. Cells were pelleted by centrifugation (4000 rpm, 10 min) (ALC, PK121R) and resuspended in 500µL of PBS. The optical densities of all suspensions were adjusted to 0.1 OD_600_. Aliquots of 180 µL of cell suspension were transferred to a black flat-bottomed 96-well plate (Greiner Bio-one, Stonehouse, UK). Eight technical replicates, for each strain tested, were in each column of the plate. The plate was transferred to a POLARstar spectrophotometer (BMG Labtech) and incubated at 37°C. Hoescht dye (H33342, 250 µM in water) was added to bacterial suspension of the plate using the plate-reader’s auto-injector to give a final concentration of 25 µM per well. Excitation and emission filters were set at 355 nm and 460 nm respectively. Readings were taken in intervals (cycles) separated by 150 s. 31 cycles were run in total. A gain multiplier of 1300 was used. Results were expressed as absolute values of fluorescence versus time.

### Proteomics

500 µL of an overnight NB culture were transferred to 50 mL NB and cells were grown at 37°C to 0.6 OD_600_. Cells were pelleted by centrifugation (10 min, 4,000 × *g*, 4°C) and resuspended in 20 mL of 30 mM Tris-HCl, pH 8 and broken by sonication using a cycle of 1 s on, 0.5 s off for 3 min at amplitude of 63% using a Sonics Vibracell VC-505TM (Sonics and Materials Inc., Newton, Connecticut, USA). The sonicated samples were centrifuged at 8,000 rpm (Sorval RC5B PLUS using an SS-34 rotor) for 15 min at 4°C to pellet intact cells and large cell debris; For envelope preparations, the supernatant was subjected to centrifugation at 20,000 rpm for 60 min at 4°C using the above rotor to pellet total envelopes. To isolate total envelope proteins, this total envelope pellet was solubilised using 200 μL of 30 mM Tris-HCl pH 8 containing 0.5% (w/v) SDS.

Protein concentrations in all samples were quantified using Biorad Protein Assay Dye Reagent Concentrate according to the manufacturer’s instructions. Proteins (5 µg/lane for envelope protein analysis) were separated by SDS-PAGE using 11% acrylamide, 0.5% bis-acrylamide (Biorad) gels and a Biorad Min-Protein Tetracell chamber model 3000X1. Gels were resolved at 200 V until the dye front had moved approximately 1 cm into the separating gel. Proteins in all gels were stained with Instant Blue (Expedeon) for 20 min and de-stained in water.

The 1 cm of gel lane was subjected to in-gel tryptic digestion using a DigestPro automated digestion unit (Intavis Ltd). The resulting peptides from each gel fragment were fractionated separately using an Ultimate 3000 nanoHPLC system in line with an LTQ-Orbitrap Velos mass spectrometer (Thermo Scientific). In brief, peptides in 1% (v/v) formic acid were injected onto an Acclaim PepMap C18 nano-trap column (Thermo Scientific). After washing with 0.5% (v/v) acetonitrile plus 0.1% (v/v) formic acid, peptides were resolved on a 250 mm × 75 μm Acclaim PepMap C18 reverse phase analytical column (Thermo Scientific) over a 150 min organic gradient, using 7 gradient segments (1-6% solvent B over 1 min, 6-15% B over 58 min, 15-32% B over 58 min, 32-40% B over 5 min, 40-90% B over 1 min, held at 90% B for 6 min and then reduced to 1% B over 1 min) with a flow rate of 300 nL/min.

Solvent A was 0.1% formic acid and Solvent B was aqueous 80% acetonitrile in 0.1% formic acid. Peptides were ionized by nano-electrospray ionization MS at 2.1 kV using a stainless-steel emitter with an internal diameter of 30 μm (Thermo Scientific) and a capillary temperature of 250°C. Tandem mass spectra were acquired using an LTQ-Orbitrap Velos mass spectrometer controlled by Xcalibur 2.1 software (Thermo Scientific) and operated in data-dependent acquisition mode. The Orbitrap was set to analyze the survey scans at 60,000 resolution (at m/z 400) in the mass range m/z 300 to 2000 and the top twenty multiply charged ions in each duty cycle selected for MS/MS in the LTQ linear ion trap. Charge state filtering, where unassigned precursor ions were not selected for fragmentation, and dynamic exclusion (repeat count, 1; repeat duration, 30 s; exclusion list size, 500) were used. Fragmentation conditions in the LTQ were as follows: normalized collision energy, 40%; activation q, 0.25; activation time 10 ms; and minimum ion selection intensity, 500 counts.

The raw data files were processed and quantified using Proteome Discoverer software v1.4 (Thermo Scientific) and searched against the UniProt *S. maltophilia* strain K279a database (4365 protein entries; UniProt accession UP000008840) using the SEQUEST (Ver. 28 Rev. 13) algorithm. Peptide precursor mass tolerance was set at 10 ppm, and MS/MS tolerance was set at 0.8 Da. Search criteria included carbamidomethylation of cysteine (+57.0214) as a fixed modification and oxidation of methionine (+15.9949) as a variable modification. Searches were performed with full tryptic digestion and a maximum of 1 missed cleavage was allowed. The reverse database search option was enabled and all peptide data was filtered to satisfy false discovery rate (FDR) of 5 %. The Proteome Discoverer software generates a reverse “decoy” database from the same protein database used for the analysis and any peptides passing the initial filtering parameters that were derived from this decoy database are defined as false positive identifications. The minimum cross-correlation factor filter was readjusted for each individual charge state separately to optimally meet the predetermined target FDR of 5 % based on the number of random false positive matches from the reverse decoy database. Thus, each data set has its own passing parameters. Protein abundance measurements were calculated from peptide peak areas using the Top 3 method (Silva *et al*., 2006) and proteins with fewer than three peptides identified were excluded. The proteomic analysis was repeated three times for each parent and mutant strain, each using a separate batch of cells. Data analysis was as follows: all raw protein abundance data were uploaded into Microsoft Excel. Raw data from each sample were normalised by division by the average abundance of all 30S and 50S ribosomal protein in that sample. A one-tailed, unpaired T-Test was used to calculate the significance of any difference in normalised protein abundance data in the three sets of data from the parent strains versus the three sets of data from the mutant derivative. A p-value of <0.05 was considered significant. The fold change in abundance for each protein in the mutant compared to its parent was calculated using the averages of normalised protein abundance data for the three biological replicates for each strain.

### Whole genome sequencing to Identify mutations

Whole genome resequencing was performed by MicrobesNG (Birmingham, UK) on a HiSeq 2500 instrument (Illumina, San Diego, CA, USA). Reads were trimmed using Trimmomatic (Bolger *et al*., 2014) and assembled into contigs using SPAdes 3.10.1 (http://cab.spbu.ru/software/spades/). Assembled contigs were mapped to reference genome for *S. maltophilia* K279a (Crossman *et al*., 2008) obtained from GenBank (accession number NC_010943) using progressive Mauve alignment software (Darling *et al*., 2010).

Mutations were checked by PCR using Phusion High Fidelity DNA Polymerase (New England Biolabs). To generate template DNA, a bacterial colony was resuspended in 100 µL of molecular biology grade water and heated at 100°C for 5 min. The sample was centrifuged at 13000 rpm for 5 min. PCR reactions were set up using 5 µL of 5X Phusion GC Buffer, 0.5 µL of dNTPs (10 mM), 1.25 µL of forward primer (10 µM), 1.25 µL of reverse primer (10 µM), 0.75 µL of DMSO, 0.25 µL of Phusion DNA Polymerase, 1 µL of DNA template, and 15 µL of molecular biology grade water. The cycling conditions were the following: 1 cycle of 98°C for 30 s, 30 cycles of: 98°C for 10 s, 62°C for 30 s, and 72°C for 30 s, 1 cycle of 72°C for 10 min for final extension.

The primers used were: *smlt0009* F 5’-GTGTGAAGAACCAGGCTGATGCCA-3’ and *smlt0009* R 5’-AGGGTGTAGCTAAGCTAAACAAT-3’. PCR products were purified using the QIAquick PCR Purification Kit (Qiagen) according to the manufacturer’s instructions. DNA concentration of purified samples was quantified using NanoDrop Lite spectrophotomer (Thermo Scientific). PCR products were sequenced by Eurofins. Sequences obtained were analysed with ClustalW OMEGA or MultiAlignPro. Alignments were represented using ESPript 3.0.

### Insertional inactivation of smlt0009

The K279a Δ*smlt0009* mutant was constructed by gene inactivation mediated by the pKNOCK suicide plasmid (Alexeyev, 1999). The *smlt0009* DNA fragment was amplified with Phusion High-Fidelity DNA Polymerase (NEB, UK) from *S. maltophilia* genomic DNA by using primers *smlt0009* KO FW (5′-GTGAAGAATCTGTCGCCGC-3′) and *smlt0009* KO RV (5′-GGATCACTTCGCCCTGGATA-3′). The PCR product was ligated into the pKNOCK-GM at SmaI site. The recombinant plasmid was then transferred into wild-type *S. maltophilia* cells by conjugation. The mutant was selected for gentamicin resistance and the mutation was confirmed by PCR using primers *smlt0009* full length FW (5’-AAAGAATTCAGTAGGAATAACGCCTGAATGC-3’) and *smlt0009* full length RV (5’-AAAGAATTCTGACGCTTACCTTTGTTGTGTG-3’).

## Acknowledgments

This work was funded by grant MR/N013646/1 to M.B.A. and K.J.H and grant MR/S004769/1 to M.B.A. from the Antimicrobial Resistance Cross Council Initiative supported by the seven United Kingdom research councils and the National Institute for Health Research. Genome sequencing was provided by MicrobesNG (http://www.microbesng.uk/), which is supported by the BBSRC (grant number BB/L024209/1). K.C. received a postgraduate scholarship from SENESCYT, Ecuador. P. D. received a scholarship from the Thai Royal Golden Jubilee Ph.D. Programme (Grant NO: PHD/0036/2558).

