## Supplementary Data for "TonB dependent uptake of β-lactam antibiotics in the opportunistic human pathogen *Stenotrophomonas maltophilia*"

**Table S1** **Normalised proteomics data for mutants M1 and M52 relative to K279a parent**

Abundance changes significantly according to a T-Test (P<0.05). Proteins that are >1.5 fold up and down regulated are highlighted green or red, respectively. Proteins are ordered based on Uniprot Accession number. Consecutive numbers are suggestive of operons. Where proteins are not detectable in one sample, it is not possible to give an accurate fold change so >100 or <0.01 are used to represent the actual change, which must be considerable.

| **Accession** | **Description** |  | **T-Test**  **_K279a vs M1_** | **Fold**  **_K279a vs M1_** | **T-Test**  **_K279a vs M52_** | **Fold**  **_K279a vs M52_** |
| --- | --- | --- | --- | --- | --- | --- |
| B2FHA2 | Putative ferredoxin oxidoreductase | Smlt0136 | 0.034 | 0.42 | 0.022 | 0.17 |
| B2FHA8 | Putative phospholipase | Smlt0142 | 0.018 | 0.53 | 0.022 | 0.56 |
| B2FHB5 | Uncharacterized protein | Smlt0149 | 0.019 | 0.53 | 0.011 | 0.53 |
| B2FHB7 | Glutamine synthetase | glnA | <0.05 | >100 | <0.05 | >100 |
| B2FHD6 | Uncharacterized protein | Smlt0170 | 0.044 | 0.62 | 0.047 | 0.61 |
| B2FHF0 | Uncharacterized protein | Smlt0184 | 0.034 | 0.58 | 0.007 | 0.46 |
| B2FHF9 | Uncharacterized protein | Smlt0193 | <0.05 | >100 | <0.05 | >100 |
| B2FHH2 | Putative TonB dependent siderophore receptor | Smlt1426 | 0 | 13.73 | 0 | 22.44 |
| B2FHH7 | Putative peptidase | Smlt1431 | 0.004 | 0.32 | 0.003 | 0.3 |
| B2FHJ2 | Putative outer membrane protein | Smlt1446 | 0.001 | 0.3 | 0.001 | 0.23 |
| B2FHJ5 | Putative phosphodiesterase-nucleotide pyrophosphatase | Smlt1449 | 0.014 | 0.47 | 0.005 | 0.41 |
| B2FHL6 | Putative ABC transporter | Smlt1471 | 0.007 | 0.63 | 0.05 | 0.62 |
| B2FHL9 | Uncharacterized protein | Smlt1474 | 0.023 | 0.54 | 0.014 | 0.51 |
| B2FHM1 | Putative transmembrane protein | Smlt1476 | 0.01 | 0.55 | 0.018 | 0.59 |
| B2FHN2 | Uncharacterized protein | Smlt1491 | 0.01 | 0.37 | 0.013 | 0.46 |
| B2FHQ4 | Putative hydrolase | Smlt2820 | <0.05 | >100 | <0.05 | >100 |
| B2FHT4 | Putative TonB dependent extracellular heme-binding protein | Smlt2850 | <0.05 | >100 | <0.05 | >100 |
| B2FHT9 | Putative iron transporter | Smlt2858 | 0.003 | 3.49 | 0.001 | 4.77 |
| B2FHX5 | Putative CBS domain protein | Smlt4098 | <0.05 | >100 | <0.05 | >100 |
| B2FI00 | Putative outer membrane Omp family protein | Smlt4123 | 0.046 | 0.6 | 0.012 | 0.49 |
| B2FI12 | Putative colicin I receptor | cirA | 0.001 | 27.93 | 0 | 76.72 |
| B2FI43 | Putative peptidyl-dipeptidase | dcp | 0.032 | 0.58 | 0.037 | 0.58 |
| B2FIA8 | Elongation factor Ts | tsf | 0.004 | 1.68 | 0.033 | 1.81 |
| B2FIA9 | 30S ribosomal protein S2 | rpsB | 0.042 | 0.71 | 0.015 | 0.65 |
| B2FIF8 | Peptidyl-prolyl cis-trans isomerase | Smlt1559 | 0.013 | 1.86 | 0.001 | 1.84 |
| B2FII5 | Putative CDP-diacylglycerol pyrophosphatase | Smlt2903 | 0.012 | 0.4 | 0.006 | 0.39 |
| B2FIJ2 | Putative L-lactate permease | lctP | 0.048 | 1.7 | 0.013 | 2 |
| B2FIL8 | Putative TonB dependent receptor protein | Smlt2937 | <0.05 | >100 | <0.05 | >100 |
| B2FIN7 | Putative TonB dependent receptor | Smlt4151 | <0.05 | 1.22 | <0.05 | >100 |
| B2FIQ3 | Putative lipid A biosynthesis lauroyl acyltransferase | htrB | 0.035 | 0.62 | 0.05 | 0.69 |
| B2FIS6 | Protease 4 | sppA | 0.011 | 0.54 | 0.006 | 0.51 |
| B2FIU9 | 60 kDa chaperonin | groL | 0.045 | 1.7 | 0.011 | 1.95 |
| B2FIV0 | 10 kDa chaperonin | groS | 0.001 | 2.49 | 0.006 | 2.92 |
| B2FJ60 | Putative transmembrane PepSY domain protein | Smlt1566 | <0.05 | >100 | <0.05 | >100 |
| B2FJ75 | Putrescine-binding periplasmic protein | potF | 0.018 | 0.54 | 0.019 | 0.58 |
| B2FJ77 | Polyamine-transporting ATPase | potG | 0.021 | 0.44 | 0.019 | 0.46 |
| B2FJB1 | Putative oar family adhesion protein | Smlt1619 | 0.032 | 0.6 | 0.023 | 0.54 |
| B2FJJ3 | Putative hydroxamate-type ferrisiderophore receptor | Smlt3022 | 0.007 | 12.51 | 0.002 | 22.63 |
| B2FJR9 | Uncharacterized protein | Smlt4275 | 0.045 | 1.59 | 0.027 | 2.19 |
| B2FJS3 | Putative HlyD family efflux protein | Smlt4279 | 0.022 | 0.54 | 0.013 | 0.47 |
| B2FJU4 | 50S ribosomal protein L13 | rplM | 0.044 | 1.25 | 0.004 | 1.43 |
| B2FJV0 | Uncharacterized protein | Smlt0387 | 0.014 | 0.54 | 0.002 | 0.4 |
| B2FK68 | Putative glutamate synthase | Smlt1693 | 0.005 | 0.38 | 0.002 | 0.28 |
| B2FK88 | Enolase | eno | <0.05 | 6.86 | <0.05 | 11.58 |
| B2FKE9 | Putative P-protein | pheA | 0.022 | 0.08 | 0.005 | 0.07 |
| B2FKJ7 | 50S ribosomal protein L9 | rplI | 0.023 | 1.51 | 0.015 | 1.62 |
| B2FKJ9 | 30S ribosomal protein S6 | rpsF | 0.003 | 1.77 | 0.009 | 1.67 |
| B2FKL0 | Putative haloacid dehalogenase hydrolase | Smlt4308 | 0.05 | 0.57 | 0.044 | 0.58 |
| B2FKM2 | Putative K(+)/H(+) antiporter subunit A/B | phaAB | 0.021 | 1.59 | 0.028 | 1.55 |
| B2FKX4 | Uncharacterized protein | Smlt0503 | <0.05 | >100 | <0.05 | >100 |
| B2FL08 | Putative transmembrane anchor protein | Smlt0538 | 0.002 | 0.25 | 0.003 | 0.34 |
| B2FL10 | Uncharacterized protein | Smlt0540 | 0.002 | 0.26 | 0.002 | 0.31 |
| B2FL33 | Alanine--tRNA ligase | alaS | <0.05 | 0.9 | <0.05 | 0.68 |
| B2FL51 | Putative ferric siderophore receptor protein | Smlt1762 | <0.05 | >100 | <0.05 | >100 |
| B2FL84 | Putative succinate dehydrogenase cytochrome b-556 subunit | sdhC | 0.019 | 0.63 | 0.005 | 0.48 |
| B2FL86 | Putative succinate dehydrogenase flavoprotein subunit | sdhA | 0.02 | 0.44 | 0.006 | 0.28 |
| B2FL87 | Succinate dehydrogenase iron-sulfur subunit | sdhB | 0.021 | 0.54 | 0.004 | 0.36 |
| B2FLB8 | Peptidyl-prolyl cis-trans isomerase | Smlt3182 | 0.038 | 0.62 | 0.007 | 0.44 |
| B2FLD3 | Dihydrolipoyl dehydrogenase | odhL | 0.033 | 1.95 | 0.017 | 2.31 |
| B2FLE4 | Putative outer membrane antigen protein | Smlt3210 | 0.023 | 0.59 | 0.002 | 0.4 |
| B2FLE9 | Putative outer membrane antigen lipoprotein | Smlt3215 | 0.034 | 0.58 | 0.013 | 0.51 |
| B2FLG5 | Uncharacterized protein | Smlt3232 | 0.025 | 0.56 | 0.049 | 0.64 |
| B2FLQ1 | D-amino acid dehydrogenase | dadA | 0.016 | 0.4 | 0.016 | 0.41 |
| B2FLR8 | Putative vitamin B12 receptor protein | Smlt0585 | 0.023 | 0.57 | 0.018 | 0.57 |
| B2FLT1 | Asparagine synthetase | Smlt0598 | <0.05 | >100 | <0.05 | >100 |
| B2FLX9 | Putative outer membrane lipoprotein | Smlt1826 | 0.03 | 0.57 | 0.01 | 0.49 |
| B2FM92 | Ribonuclease E | rnE | 0.003 | 2.12 | 0.01 | 1.92 |
| B2FME8 | Putative DNA binding protein/regulator | Smlt3308 | <0.05 | >100 | <0.05 | >100 |
| B2FMF2 | Uncharacterized protein | Smlt3312 | 0.04 | 0.57 | 0.018 | 0.51 |
| B2FMI1 | Putative transmembrane protein | Smlt4498 | 0.025 | 0.57 | 0.01 | 0.55 |
| B2FMN3 | Putative transmembrane anchor short-chain dehydrogenase | Smlt0632 | 0.03 | 0.62 | 0.035 | 0.68 |
| B2FMP6 | Putative electron transfer flavoprotein subunit beta | etfS | 0.034 | 2.22 | 0.001 | 2.39 |
| B2FMY6 | Chaperone protein DnaJ | dnaJ | 0.022 | 1.66 | 0.028 | 3.19 |
| B2FN04 | Putative reductase Smlt2015 | Smlt2015 | <0.05 | 2.18 | <0.05 | 2.38 |
| B2FN37 | Putative N-acetylmuramoyl-L-alanine amidase | Smlt3330 | 0.038 | 0.56 | 0.008 | 0.41 |
| B2FN47 | Putative TonB dependent receptor | Smlt3340 | 0.006 | 0.37 | 0.003 | 0.37 |
| B2FNC5 | Putative dipeptidyl peptidase | Smlt4581 | 0.006 | 0.48 | 0.014 | 0.53 |
| B2FND7 | Uncharacterized protein | Smlt4593 | 0.003 | 0.37 | 0.003 | 0.45 |
| B2FNF4 | Putative glycosyl transferase | Smlt4611 | 0.028 | 0.53 | 0.009 | 0.46 |
| B2FNG4 | Putative Major Facilitator Superfamily transmembrane transport protein | Smlt4621 | <0.05 | >100 | <0.05 | >100 |
| B2FNG5 | Conserved hypothetical exported protein | Smlt4622 | <0.05 | 5.89 | <0.05 | 5.35 |
| B2FNG6 | Acetyl-coenzyme A synthetase | acsA | <0.05 | >100 | <0.05 | >100 |
| B2FNP5 | 30S ribosomal protein S1 | rpsA | 0.01 | 1.72 | 0.005 | 1.84 |
| B2FNQ0 | Putative epimerase/dehydratase polysaccharide-related biosynthesis protein | wbiI | 0.012 | 0.48 | 0.027 | 0.56 |
| B2FNX1 | NADH-quinone oxidoreductase subunit I | nuoI | 0.018 | 0.48 | 0.003 | 0.31 |
| B2FNX6 | NADH-quinone oxidoreductase subunit D | nuoD | 0.018 | 0.57 | 0.004 | 0.39 |
| B2FNX7 | NADH-quinone oxidoreductase subunit C | nuoC | 0.011 | 0.51 | 0.001 | 0.33 |
| B2FNX8 | NADH-quinone oxidoreductase subunit B | nuoB | 0.02 | 0.51 | 0.003 | 0.34 |
| B2FNY0 | Seneral secretory pathway protein | secG | 0.001 | 0.43 | 0.003 | 0.52 |
| B2FNY4 | Putative lipopolysaccharide core biosynthesis glycosyl transferase | Smlt3411 | 0.04 | 0.64 | 0.046 | 0.67 |
| B2FP19 | Putative TonB dependent receptor protein | Smlt3449 | 0.039 | 1.64 | 0.01 | 2.53 |
| B2FP56 | Ferrochelatase | hemH | 0.048 | 0.5 | 0.028 | 0.44 |
| B2FP85 | Putative ABC transporter toluene tolerance exported protein | Smlt4673 | 0.005 | 1.75 | 0.004 | 1.76 |
| B2FPA5 | Membrane protein insertase YidC | yidC | 0.028 | 0.56 | 0.014 | 0.52 |
| B2FPD1 | Putative glycosyl transferase | Smlt0784 | 0.011 | 0.45 | 0.015 | 0.54 |
| B2FPD9 | Probable Na+ dependent nucleoside transporter | Smlt0792 | 0.003 | 0.73 | 0.032 | 0.76 |
| B2FPE2 | Putative exported heme receptor protein | huvA | 0 | 22.44 | 0 | 43.14 |
| B2FPE4 | Uncharacterized protein | Smlt0797 | <0.05 | >100 | <0.05 | >100 |
| B2FPF2 | Putative transmembrane protein | Smlt0805 | 0.044 | 0.6 | 0.014 | 0.49 |
| B2FPF9 | Prolipoprotein diacylglyceryl transferase | lgt | 0.005 | 0.44 | 0.01 | 0.47 |
| B2FPN1 | Putative TonB dependent receptor | Smlt2175 | 0.009 | 2.98 | 0.013 | 2.65 |
| B2FPQ9 | Putative TonB domain protein | Smlt3477 | <0.05 | >100 | <0.05 | >100 |
| B2FPR0 | Putative TonB dependent receptor | Smlt3478 | 0.019 | 3.83 | 0.014 | 4.26 |
| B2FPS0 | Putative exported LysM bacterial cell wall related protein | Smlt3488 | 0.02 | 0.49 | 0.009 | 0.43 |
| B2FPS4 | Putative transmembrane protein | Smlt3492 | 0.008 | 0.47 | 0.012 | 0.53 |
| B2FPV6 | Putative thiolase | Smlt3525 | <0.05 | 0.38 | <0.05 | 0.62 |
| B2FPY0 | Uncharacterized protein | Smlt3550 | 0.01 | 0.51 | 0.04 | 0.66 |
| B2FPY7 | Putative alkyl hydroperoxide reductase subunit c | ahpC | 0.024 | 3.37 | 0.004 | 4.5 |
| B2FQ28 | Putative TonB dependent receptor protein | Smlt0885 | 0.016 | 0.54 | 0.032 | 0.59 |
| B2FQ38 | DNA-directed RNA polymerase subunit beta | rpoB | 0.037 | 2.71 | 0.018 | 3 |
| B2FQ39 | DNA-directed RNA polymerase subunit beta' | rpoC | 0.033 | 2.47 | 0.022 | 2.28 |
| B2FQ42 | Elongation factor G | fusA | 0.046 | 2.16 | 0.009 | 1.85 |
| B2FQ45 | 50S ribosomal protein L3 | rplC | 0.006 | 1.37 | 0.031 | 1.28 |
| B2FQ48 | 50S ribosomal protein L2 | rplB | 0.037 | 1.57 | 0.04 | 1.4 |
| B2FQ50 | 50S ribosomal protein L22 | rplV | 0.019 | 1.77 | 0.025 | 2.14 |
| B2FQD3 | Putative carbon starvation protein A | cstA | 0 | 3.8 | 0.008 | 3.55 |
| B2FQJ8 | 30S ribosomal protein S8 | rpsH | 0.028 | 0.71 | 0.031 | 0.73 |
| B2FQK6 | 30S ribosomal protein S11 | rpsK | 0.029 | 1.37 | 0 | 1.73 |
| B2FQK8 | DNA-directed RNA polymerase subunit alpha | rpoA | 0.043 | 2.11 | 0.016 | 1.9 |
| B2FQL8 | Malate dehydrogenase | mdh | 0.043 | 2.36 | 0.034 | 7.46 |
| B2FQN3 | Uncharacterized protein | Smlt0960 | 0.008 | 0.43 | 0.007 | 0.43 |
| B2FQN4 | L-threonine 3-dehydrogenase | tdh | <0.05 | >100 | <0.05 | >100 |
| B2FR08 | Putative TonB dependent receptor | Smlt3645 | 0.003 | 2.73 | 0.007 | 3.67 |
| B2FR62 | Putative autotransporter | Smlt1001 | 0.001 | 0.1 | 0 | 0.09 |
| B2FRC4 | Putative TonB-dependent receptor | Smlt1067 | 0.003 | 4.64 | 0.003 | 4.28 |
| B2FRE3 | Putative esterase | Smlt2353 | <0.05 | >100 | <0.05 | >100 |
| **B2FRE4** | **Putative ATP-binding protein** | **Smlt2354** | <0.05 | >100 | <0.05 | >100 |
| **B2FRE5** | **Putative binding-protein-dependent transport lipoprotein** | **Smlt2355** | 0 | 14.81 | 0 | 28.03 |
| **B2FRE6** | **Putative FecCD-family transmembrane transport protein** | **Smlt2356** | <0.05 | >100 | <0.05 | >100 |
| **B2FRE7** | **Hemin import ATP-binding protein HmuV** | **hmuV** | <0.05 | >100 | <0.05 | >100 |
| B2FRZ9 | Putative iron transport receptor protein | Smlt1148 | 0.01 | 2.17 | 0.005 | 2.57 |
| B2FS15 | Putative histone-like protein | Smlt1164 | 0.02 | 3.04 | 0.001 | 5.72 |
| B2FSE3 | Putative HlyD-family secretion protein | smeM | 0.027 | 0.65 | 0.001 | 0.39 |
| B2FSE4 | Putative TonB dependent receptor | Smlt3789 | 0.002 | 5.18 | 0 | 4.28 |
| B2FSE9 | Conserved hypothetical exported protein | Smlt3796 | 0.002 | 0.34 | 0.001 | 0.34 |
| B2FSF0 | Fructose-bisphosphate aldolase | alf1 | 0.001 | 2.16 | 0 | 2.92 |
| B2FSF6 | Glyceraldehyde-3-phosphate dehydrogenase | gap | 0.009 | 2.79 | 0 | 2.91 |
| B2FSQ7 | Putative TonB-dependent receptor | Smlt1233 | 0 | 20.43 | 0 | 45.94 |
| B2FSS6 | Putative PTS system, fructose-specific IIBC component | fruA | 0.046 | 1.65 | 0.034 | 1.8 |
| B2FST2 | Putative TonB dependent receptor | Smlt2566 | 0.019 | 2.9 | 0.003 | 3.07 |
| B2FT59 | Putative extracellular heme-binding protein | Smlt3898 | 0.006 | 3.3 | 0.001 | 8.38 |
| B2FT68 | Putative transcriptional regulator | Smlt3907 | 0.015 | 0.5 | 0.016 | 0.53 |
| B2FT80 | DNA polymerase III subunit beta | dnaN | <0.05 | >100 | <0.05 | >100 |
| B2FT86 | Conserved hypothetical TPR repeat family protein | Smlt0008 | 0.001 | 2.82 | 0 | 3.24 |
| **B2FT87** | **Putative proline-rich TonB dependent receptor protein** | **Smlt0009** | 0.045 | 0.53 | 0.015 | 0.47 |
| B2FTK5 | Putative ferric siderophore receptor | Smlt2650 | <0.05 | 7.93 | <0.05 | 12.31 |
| B2FTR3 | Conserved hypothetical exported protein | Smlt2712 | <0.05 | >100 | <0.05 | >100 |
| B2FTR4 | Conserved hypothetical exported protein | Smlt2713 | <0.05 | >100 | <0.05 | >100 |
| B2FTR5 | Putative TonB dependent protein, possible siderophore receptor | Smlt2714 | <0.05 | >100 | <0.05 | >100 |
| B2FTS4 | Uncharacterized protein | pcm | 0.025 | 0.64 | 0.01 | 0.49 |
| B2FTS5 | Protein CyaE | tolC | 0.001 | 0.34 | 0 | 0.3 |
| B2FTS8 | Putative lipid A biosynthesis lauroyl acyltransferase | htrB | 0.045 | 0.57 | 0.03 | 0.55 |
| B2FTY9 | Putative endonuclease/exonuclease/phosphatase family protein | Smlt3995 | 0.012 | 0.47 | 0.006 | 0.37 |
| B2FU12 | Putative iron transporter protein | Smlt0049 | 0.015 | 6.19 | 0.013 | 3.74 |
| B2FU42 | Putative TonB dependent receptor protein | Smlt0083 | 0.009 | 0.45 | 0.003 | 0.35 |
| B2FU43 | Acid phosphatase | Smlt0084 | 0 | 0.45 | 0 | 0.39 |
| B2FU57 | Putative lipoprotein E (Outer membrane protein p4) | hel | 0.029 | 0.53 | 0.041 | 0.61 |
| B2FUD9 | Putative HlyD family secretion protein | Smlt1406 | 0.033 | 0.57 | 0.039 | 0.63 |
| B2FUL1 | Putative respiratory nitrate reductase subunit | narH | 0.04 | 0.45 | 0.018 | 0.38 |
| B2FUN9 | Putative TonB-dependent receptor | fhuE | 0.001 | 10.25 | 0 | 16.63 |
| B2FUR1 | Putative TonB-dependent outer membrane receptor protein | Smlt4026 | 0.05 | 0.64 | 0.007 | 0.37 |
| B2FUV1 | Putative multidrug resistance outer membrane protein | smeF | 0.042 | 0.65 | 0.009 | 0.54 |

**Table S2 Normalised proteomics data for K279a Δsmlt0009 relative to K279a parent**

Abundance changes significantly according to a T-Test (P<0.05). Proteins that are >1.5 fold up and down regulated are highlighted green or red, respectively. Proteins are ordered based on Uniprot Accession number. Consecutive numbers are suggestive of operons. Where proteins are not detectable in one sample, it is not possible to give an accurate fold change so >100 or <0.01 are used to represent the actual change, which must be considerable.

| Accession | Description |  | T-Test | Fold |
| --- | --- | --- | --- | --- |
|  |  |  | K279a vs  K279a *Δsmlt0009* | K279a vs  K279a *Δsmlt0009* |
| B2FH97 | Putative modulator of DNA gyrase | Smlt0130 | 0.000 | <0.01 |
| B2FHA0 | Putative survival protein HipA | hipA | 0.004 | <0.01 |
| B2FHA2 | Putative ferredoxin oxidoreductase | Smlt0136 | 0.000 | <0.01 |
| B2FHA8 | Putative phospholipase | Smlt0142 | 0.002 | <0.01 |
| B2FHB0 | Putative transmembrane acetyltransferase | Smlt0144 | 0.005 | 7.51 |
| B2FHB1 | Uncharacterized protein | Smlt0145 | 0.009 | <0.01 |
| B2FHB2 | Putative TPR repeat protein | Smlt0146 | 0.016 | 0.46 |
| B2FHB5 | Uncharacterized protein | Smlt0149 | 0.000 | 0.03 |
| B2FHB7 | Glutamine synthetase | glnA | 0.000 | <0.01 |
| B2FHC6 | Superoxide dismutase [Cu-Zn] | sodC1 | 0.008 | 2.51 |
| B2FHC7 | Superoxide dismutase [Cu-Zn] | sodC2 | 0.007 | 2.78 |
| B2FHD0 | Putative 3-ketoacyl-CoA thiolase | fadI | 0.002 | 2.56 |
| B2FHD1 | Putative transmembrane HemY porphyrin biosynthesis protein | Smlt0165 | 0.000 | 4.40 |
| B2FHD6 | Uncharacterized protein | Smlt0170 | 0.015 | 2.00 |
| B2FHE8 | Putative transmembrane protein | Smlt0182 | 0.001 | 12.95 |
| B2FHF0 | Uncharacterized protein | Smlt0184 | 0.005 | 5.79 |
| B2FHF3 | Putative peroxidase | Smlt0187 | 0.000 | <0.01 |
| B2FHF6 | Putative TonB dependent receptor protein | Smlt0190 | 0.030 | 1.67 |
| B2FHG4 | Putative electron transfer flavoprotein-ubiquinone oxidoreductase | Smlt0198 | 0.015 | 2.03 |
| B2FHG5 | Putative glutaryl-CoA dehydrogenase | GCDH | 0.000 | <0.01 |
| B2FHG9 | Uncharacterized protein | Smlt0204 | 0.007 | 3.79 |
| B2FHH0 | Putative 3-oxoacyl-[acyl-carrier-protein] synthase 3 | fabH | 0.000 | <0.01 |
| B2FHH2 | Putative TonB dependent siderophore receptor | Smlt1426 | 0.001 | 54.02 |
| B2FHI0 | Putative acyl-CoA synthetase | Smlt1434 | 0.016 | 1.58 |
| B2FHJ2 | Putative outer membrane protein | Smlt1446 | 0.050 | 0.24 |
| B2FHJ5 | Putative phosphodiesterase-nucleotide pyrophosphatase | Smlt1449 | 0.023 | 1.87 |
| B2FHK0 | Uncharacterized protein | Smlt1454 | 0.005 | 8.89 |
| B2FHK4 | Putative monooxygenase | Smlt1459 | 0.001 | <0.01 |
| B2FHK6 | ATP-dependent RNA helicase DeaD | deaD | 0.004 | 0.24 |
| B2FHL0 | Putative GGDEF and GAF signalling protein | Smlt1465 | 0.000 | <0.01 |
| B2FHM1 | Putative transmembrane protein | Smlt1476 | 0.000 | 2.19 |
| B2FHM8 | Putative PHB depolymerase | Smlt1486 | 0.000 | <0.01 |
| B2FHN1 | Acetyl-coenzyme A carboxylase carboxyl transferase subunit alpha | accA | 0.015 | 0.39 |
| B2FHN3 | DNA-directed DNA polymerase | dnaE | 0.043 | 0.47 |
| B2FHN5 | Lipid-A-disaccharide synthase | lpxB | 0.002 | 3.54 |
| B2FHR6 | Putative Fe/S OXIDOREDUCTASE | Smlt2832 | 0.000 | <0.01 |
| B2FHR9 | Putative TonB-dependent outer membrane protein | Smlt2835 | 0.003 | 4.76 |
| B2FHT9 | Putative iron transporter | Smlt2858 | 0.003 | 4.92 |
| B2FHX5 | Putative CBS domain protein | Smlt4098 | 0.000 | 8.61 |
| B2FHY2 | Putative two-component response regulator transcriptional regulatory protein | Smlt4105 | 0.000 | 0.09 |
| B2FHY3 | Histidine kinase | Smlt4106 | 0.000 | 1.91 |
| B2FHY5 | Bifunctional protein GlmU | glmU | 0.004 | <0.01 |
| B2FHY8 | ATP synthase subunit beta | atpD | 0.006 | 2.16 |
| B2FHY9 | ATP synthase gamma chain | atpG | 0.017 | 2.52 |
| B2FHZ0 | ATP synthase subunit alpha | atpA | 0.006 | 2.67 |
| B2FHZ1 | ATP synthase subunit delta | atpH | 0.002 | 2.51 |
| B2FHZ2 | ATP synthase subunit b | atpF | 0.002 | 3.42 |
| B2FHZ6 | Uncharacterized protein | Smlt4119 | 0.013 | 3.48 |
| B2FHZ7 | Putative dihydrolipoamide dehydrogenase | lpdA | 0.010 | 0.29 |
| B2FHZ8 | Acetyltransferase component of pyruvate dehydrogenase complex | pdhB | 0.031 | 0.58 |
| B2FI00 | Putative outer membrane Omp family protein | Smlt4123 | 0.009 | 3.81 |
| B2FI03 | Putative exported lipoprotein | Smlt4126 | 0.009 | 4.10 |
| B2FI05 | Conserved hypothetical exported protein | Smlt4128 | 0.002 | 0.24 |
| B2FI12 | Putative colicin I receptor | cirA | 0.003 | 62.92 |
| B2FI14 | Putative transmembrane protein | Smlt4137 | 0.019 | 1.52 |
| B2FI22 | Putative extracellular serine protease | Smlt4145 | 0.008 | <0.01 |
| B2FI26 | Putative haloalkane dehalogenase | dhaA | 0.033 | 1.93 |
| B2FI28 | Putative AMP-binding protein | Smlt0208 | 0.013 | 2.05 |
| B2FI33 | Probable protein kinase UbiB | ubiB | 0.001 | 1.89 |
| B2FI39 | Pseudouridine synthase | truC | 0.001 | <0.01 |
| B2FI42 | Putative acyltransferase | Smlt0222 | 0.003 | 1.94 |
| B2FI43 | Putative peptidyl-dipeptidase Dcp (Dipeptidyl carboxypeptidase) | dcp | 0.003 | 3.19 |
| B2FI64 | Ribonucleoside-diphosphate reductase | RRM1 | 0.007 | 0.37 |
| B2FI65 | Ribonucleoside-diphosphate reductase subunit beta | Smlt0248 | 0.030 | 0.36 |
| B2FI77 | Putative bifunctional NMN adenylyltransferase/nudix hydrolase | Smlt0261 | 0.013 | 0.35 |
| B2FI91 | tRNA-dihydrouridine synthase | dusA | 0.000 | <0.01 |
| B2FI94 | Histidine kinase | Smlt0278 | 0.048 | 1.67 |
| B2FI99 | UDP-3-O-acylglucosamine N-acyltransferase | lpxD | 0.015 | 0.23 |
| B2FIA0 | Outer membrane protein assembly factor BamA | bamA | 0.019 | 2.48 |
| B2FIA7 | Putative transmembrane GGDEF family signalling protein | Smlt1505 | 0.032 | 0.42 |
| B2FIB6 | Methionine aminopeptidase | map | 0.005 | 0.28 |
| B2FIE0 | Putative two-component system, response regulator transcriptional regulatory protein | Smlt1540 | 0.003 | 0.21 |
| B2FIE1 | Histidine kinase | Smlt1541 | 0.040 | 5.52 |
| B2FII5 | Putative CDP-diacylglycerol pyrophosphatase | Smlt2903 | 0.050 | 1.77 |
| B2FII9 | D-lactate dehydrogenase | dld | 0.001 | 1.83 |
| B2FIJ0 | L-lactate dehydrogenase | lldD | 0.001 | 7.57 |
| B2FIJ2 | Putative L-lactate permease | lctP | 0.001 | 5.73 |
| B2FIM3 | Putative orn/arg/lys decarboxylase | Smlt2942 | 0.001 | <0.01 |
| B2FIM5 | Uncharacterized protein | Smlt2944 | 0.001 | <0.01 |
| B2FIM6 | Putative thiol:disulfide interchange protein | dsbG | 0.001 | 3.92 |
| B2FIQ1 | RNA polymerase sigma factor RpoD | rpoD | 0.002 | 0.33 |
| B2FIQ3 | Putative lipid A biosynthesis lauroyl acyltransferase | htrB | 0.003 | 2.12 |
| B2FIQ4 | GTP cyclohydrolase II | Smlt4168 | 0.005 | 0.27 |
| B2FIQ7 | Putative CDP-Glycerol:Poly(Glycerophosphate) glycerophosphotransferase | Smlt4171 | 0.040 | 1.51 |
| B2FIQ8 | Putative lipopolysaccharide core biosynthesis glycosyl transferase | waaE | 0.004 | 4.30 |
| B2FIQ9 | Putative glycosyltransferase | Smlt4173 | 0.029 | 1.74 |
| B2FIR5 | Putative LysM family cell wall degradation protein | Smlt4179 | 0.003 | <0.01 |
| B2FIR8 | Putative fimbrial protein (Pilin) | Smlt4182 | 0.002 | 2.82 |
| B2FIR9 | Putative transmembrane RDD family protein | Smlt4183 | 0.002 | 2.31 |
| B2FIS6 | Protease 4 | sppA | 0.009 | 2.08 |
| B2FIS7 | Putative drug resistance transport protein | Smlt4191 | 0.000 | <0.01 |
| B2FIS9 | Putative transmembrane protein | Smlt4193 | 0.004 | 2.04 |
| B2FIT3 | Putative pyridine nucleotide-disulphide oxidoreductase | ndh | 0.000 | 4.36 |
| B2FIT9 | Phospho-2-dehydro-3-deoxyheptonate aldolase | aroG | 0.007 | <0.01 |
| B2FIU9 | 60 kDa chaperonin | groL | 0.027 | 1.74 |
| B2FIV3 | Putative transmembrane Thiol:disulfide Interchange Protein | Smlt4218 | 0.010 | 2.80 |
| B2FIX1 | Uncharacterized protein | Smlt0297 | 0.001 | <0.01 |
| B2FIX2 | Uncharacterized protein | Smlt0298 | 0.000 | <0.01 |
| B2FJ12 | Putative ankyrin repeat protein | Smlt0340 | 0.000 | <0.01 |
| B2FJ17 | Tryptophan--tRNA ligase | trpS | 0.000 | <0.01 |
| B2FJ30 | Putative TonB dependent receptor protein | Smlt0359 | 0.047 | 1.60 |
| B2FJ45 | Putative haloacid dehalogenase hydrolase | Smlt0374 | 0.000 | <0.01 |
| B2FJ51 | Uncharacterized protein | Smlt0380 | 0.001 | 3.61 |
| B2FJ52 | Putative protease | Smlt0381 | 0.023 | 2.72 |
| B2FJ63 | Uncharacterized protein | Smlt1569 | 0.001 | <0.01 |
| B2FJ68 | Bacteriohemerythrin | Smlt1574 | 0.000 | <0.01 |
| B2FJ75 | Putrescine-binding periplasmic protein | potF | 0.008 | 1.96 |
| B2FJ81 | Putative transmembrane magnesium/cobalt transport protein | corA | 0.002 | 2.58 |
| B2FJ87 | Endoribonuclease YbeY | ybeY | 0.001 | <0.01 |
| B2FJ88 | Putative PhoH-like ATP-binding protein | Smlt1596 | 0.031 | 0.41 |
| B2FJ92 | tRNA-2-methylthio-N(6)-dimethylallyladenosine synthase | miaB | 0.000 | <0.01 |
| B2FJC0 | UvrABC system protein B | uvrB | 0.013 | 0.27 |
| B2FJC8 | Histidine kinase | Smlt1636 | 0.004 | 4.90 |
| B2FJD5 | Putative transcription accessory protein | tex | 0.016 | 0.23 |
| B2FJE9 | Putative transmembrane protein | Smlt2977 | 0.014 | 2.98 |
| B2FJG0 | Conserved hypothetical exported protein | Smlt2989 | 0.011 | 2.31 |
| B2FJH4 | Putative type IV secretion system conjugal transfer protein | Smlt3003 | 0.000 | <0.01 |
| B2FJH9 | Putative transmembrane conjugative DNA transfer protein | Smlt3008 | 0.019 | 1.53 |
| B2FJJ3 | Putative hydroxamate-type ferrisiderophore receptor | Smlt3022 | 0.002 | 12.87 |
| B2FJP0 | Ribosomal protein L11 methyltransferase | prmA | 0.000 | <0.01 |
| B2FJQ5 | Putative two-component response regulator transcriptional regulatory protein | Smlt4260 | 0.029 | <0.01 |
| B2FJQ6 | Cell division protein FtsX | ftsX | 0.004 | 2.85 |
| B2FJQ7 | Putative cell division ATP-binding protein FtsE | ftsE | 0.009 | 2.07 |
| B2FJQ8 | ATP-dependent RNA helicase RhlB | rhlB | 0.003 | 0.43 |
| B2FJR0 | Transcription termination factor Rho | rho | 0.002 | 0.25 |
| B2FJR1 | Putative transmembrane GGDEF signalling protein | Smlt4266 | 0.001 | 1.72 |
| B2FJR8 | Putative isocitrate dehydrogenase [NADP] | icd | 0.017 | 2.35 |
| B2FJR9 | Uncharacterized protein | Smlt4275 | 0.001 | 5.52 |
| B2FJS3 | Putative multidrug efflux system HlyD family transmembrane protein | Smlt4279 | 0.001 | 2.75 |
| B2FJT2 | Putative transmembrane protein | Smlt4289 | 0.005 | 3.39 |
| B2FJT5 | Uncharacterized protein | Smlt4292 | 0.000 | <0.01 |
| B2FJT6 | Putative transmembrane protein | Smlt4293 | 0.008 | 2.49 |
| B2FJT8 | Putative GGDEF signaling protein | Smlt4295 | 0.015 | 0.22 |
| B2FJV5 | Putative LysR family transcriptional regulator | Smlt0393 | 0.003 | <0.01 |
| B2FJV8 | Putative oxidoreductase | Smlt0396 | 0.007 | 0.29 |
| B2FJV9 | Uncharacterized protein | Smlt0397 | 0.001 | 3.55 |
| B2FJW0 | Arginine--tRNA ligase | argS | 0.000 | <0.01 |
| B2FJW3 | Putative pantothenate biosynthesis protein | dfp | 0.013 | 0.35 |
| B2FJW5 | Putative phosphomannomutase | spgM | 0.003 | 1.71 |
| B2FJX3 | Putative endonuclease/exonuclease/phosphatase family protein | Smlt0412 | 0.001 | 0.07 |
| B2FJX8 | Tyrosine--tRNA ligase | tyrS | 0.035 | 0.27 |
| B2FJX9 | Putative aminopeptidase | Smlt0418 | 0.001 | 3.79 |
| B2FJY1 | Uncharacterized protein | Smlt0420 | 0.000 | 4.23 |
| B2FJY7 | Putative bifunctional PutA protein [includes: proline dehydrogenase delta-1-pyrroline-5-carboxylate dehydrogenase] | putA | 0.000 | 0.18 |
| B2FJZ7 | Putative cytochrome oxidase assembly protein | Smlt0436 | 0.000 | <0.01 |
| B2FK02 | DNA primase | dnaG | 0.000 | <0.01 |
| B2FK06 | tRNA N6-adenosine threonylcarbamoyltransferase | tsaD | 0.023 | 0.26 |
| B2FK09 | Putative mechanosensitive ion channel | Smlt0448 | 0.027 | 2.11 |
| B2FK11 | Uncharacterized protein | Smlt0450 | 0.000 | 7.03 |
| B2FK19 | Putative biopolymer transport protein | Smlt1638 | 0.002 | 2.93 |
| B2FK21 | Lipid A export ATP-binding/permease protein MsbA | msbA | 0.003 | 2.88 |
| B2FK22 | Tetraacyldisaccharide 4'-kinase | lpxK | 0.008 | 4.75 |
| B2FK25 | Uncharacterized protein | Smlt1644 | 0.000 | <0.01 |
| B2FK57 | Putative AraC family transcriptional regulator | Smlt1682 | 0.000 | <0.01 |
| B2FK65 | Coproporphyrinogen-III oxidase | hemN | 0.002 | <0.01 |
| B2FK71 | Peptidyl-prolyl cis-trans isomerase | fkbP | 0.010 | 3.92 |
| B2FK77 | Putative transmembrane CorC/HlyC family transporter | Smlt1704 | 0.002 | 2.76 |
| B2FK78 | Putative pit accessory protein | Smlt1705 | 0.000 | 3.01 |
| B2FK82 | DNA topoisomerase 4 subunit B | parE | 0.044 | 0.58 |
| B2FK84 | CTP synthase | pyrG | 0.014 | 0.22 |
| B2FK85 | 2-dehydro-3-deoxyphosphooctonate aldolase | kdsA | 0.000 | <0.01 |
| B2FK94 | 5'-nucleotidase SurE | surE | 0.000 | <0.01 |
| B2FK97 | Putative subfamily M23B unassigned peptidase | Smlt1724 | 0.006 | 2.13 |
| B2FKA2 | ATP-dependent zinc metalloprotease FtsH | ftsH | 0.001 | 2.93 |
| B2FKE1 | Putative LysR-family transcriptional regulator | Smlt3089 | 0.009 | 0.29 |
| B2FKE4 | Serine--tRNA ligase | serS | 0.002 | 0.13 |
| B2FKF1 | Conserved hypothetical FHA domain protein | Smlt3099 | 0.003 | 2.24 |
| B2FKI6 | DNA gyrase subunit A | gyrA | 0.001 | <0.01 |
| B2FKI7 | Methylthioribose-1-phosphate isomerase | mtnA | 0.021 | 0.23 |
| B2FKJ1 | DNA ligase | ligA | 0.005 | 0.25 |
| B2FKJ3 | Cell division protein ZipA | zipA | 0.002 | 3.21 |
| B2FKJ4 | Chromosome partition protein Smc | smc | 0.022 | 0.25 |
| B2FKK2 | Asparagine--tRNA ligase | asnS | 0.005 | 0.34 |
| B2FKK7 | S-adenosylmethionine decarboxylase proenzyme | speD | 0.009 | <0.01 |
| B2FKK8 | Putative cyclic AMP receptor protein,catabolite gene activator | cap | 0.008 | 0.48 |
| B2FKL0 | Putative haloacid dehalogenase hydrolase | Smlt4308 | 0.002 | 2.71 |
| B2FKL5 | Putative anthranilate synthase component I | trpE | 0.039 | 0.28 |
| B2FKN2 | Putative 4-hydroxyphenylpyruvate dioxygenase | Smlt4330 | 0.021 | 0.24 |
| B2FKN5 | Uncharacterized protein | Smlt4334 | 0.004 | 6.64 |
| B2FKN6 | Putative peptide transport protein | Smlt4335 | 0.032 | 3.10 |
| B2FKN9 | Putative branched-chain alpha keto acid dehydrogenase E1 beta subunit | Smlt4339 | 0.010 | 5.59 |
| B2FKR1 | Polyamine aminopropyltransferase | speE | 0.000 | <0.01 |
| B2FKR2 | Putative beta-hydroxylase | Smlt4366 | 0.001 | 4.17 |
| B2FKU8 | Putative acetyl-CoA hydrolase/transferase | Smlt0471 | 0.005 | <0.01 |
| B2FKV6 | Putative NADH dehydrogenase/NAD(P)H nitroreductase Smlt0482 | Smlt0482 | 0.008 | 0.23 |
| B2FKV7 | Conserved hypothetical exported protein | Smlt0483 | 0.014 | 3.13 |
| B2FKW2 | Pyruvate dehydrogenase E1 component | Smlt0490 | 0.005 | 0.32 |
| B2FKW6 | Uncharacterized protein | Smlt0496 | 0.000 | <0.01 |
| B2FL07 | Uncharacterized protein | Smlt0537 | 0.004 | <0.01 |
| B2FL08 | Putative transmembrane anchor protein | Smlt0538 | 0.002 | 0.34 |
| B2FL10 | Uncharacterized protein | Smlt0540 | 0.009 | 0.64 |
| B2FL20 | Putative ZINC METALLOpeptidase | Smlt0550 | 0.002 | 3.88 |
| B2FL24 | Dihydropteroate synthase | folP | 0.000 | <0.01 |
| B2FL27 | GTPase HflX | hflX | 0.005 | 0.56 |
| B2FL31 | Protein RecA | recA | 0.002 | 0.30 |
| B2FL46 | Putative cytochrome c family protein | Smlt1757 | 0.000 | <0.01 |
| B2FL47 | Putative alcohol dehydrogenase cytochrome c subunit | adhB | 0.005 | <0.01 |
| B2FL54 | Uncharacterized protein | Smlt1766 | 0.018 | 2.27 |
| B2FL62 | Conserved hypothetical exported protein | Smlt1774 | 0.011 | 3.81 |
| B2FL65 | Poly(A) polymerase I | pcnB | 0.050 | 0.43 |
| B2FL73 | Histidine kinase | Smlt1785 | 0.003 | 2.10 |
| B2FL82 | Putative ABC transporter ATP-binding subunit | Smlt1794 | 0.000 | 3.33 |
| B2FL90 | Putative lipoprotein releasing system transmembrane protein | Smlt1802 | 0.003 | 3.72 |
| B2FL91 | Lipoprotein-releasing system ATP-binding protein LolD | lolD | 0.001 | 5.78 |
| B2FL95 | Signal recognition particle receptor FtsY | ftsY | 0.009 | 0.61 |
| B2FLA1 | Conserved hypothetical exported protein | Smlt3164 | 0.002 | 4.98 |
| B2FLA2 | Putative 5-nucleotidase | Smlt3165 | 0.000 | <0.01 |
| B2FLA4 | Putative NAD-dependent glutamate dehydrogenase | Smlt3167 | 0.018 | 0.49 |
| B2FLA6 | Putative RND family acriflavine resistance protein A | smeG | 0.000 | 3.79 |
| B2FLB7 | Putative nucleotide sugar dehydrogenase | Smlt3181 | 0.006 | 0.29 |
| B2FLB8 | Peptidyl-prolyl cis-trans isomerase | Smlt3182 | 0.006 | 1.71 |
| B2FLC2 | Putative ABC transporter, ATP-binding protein | Smlt3186 | 0.001 | 2.25 |
| B2FLC3 | Putative transmembrane protein | Smlt3187 | 0.003 | 1.97 |
| B2FLC6 | Conserved hypothetical exported protein with low similarity to flagellar FliL protein | Smlt3190 | 0.006 | 2.90 |
| B2FLC8 | Adenylosuccinate lyase | purB | 0.050 | 0.44 |
| B2FLC9 | Uncharacterized protein | Smlt3195 | 0.024 | 0.40 |
| B2FLD2 | Dihydrolipoyllysine-residue succinyltransferase component of 2-oxoglutarate dehydrogenase complex | sucB | 0.012 | 0.40 |
| B2FLD3 | Dihydrolipoyl dehydrogenase | odhL | 0.016 | 0.51 |
| B2FLD8 | Putative replicative DNA helicase | dnaB | 0.003 | 0.25 |
| B2FLE4 | Putative outer membrane antigen protein | Smlt3210 | 0.002 | 7.82 |
| B2FLE7 | Putative ThiF domain protein | Smlt3213 | 0.000 | <0.01 |
| B2FLE9 | Putative outer membrane antigen lipoprotein | Smlt3215 | 0.008 | 4.34 |
| B2FLF6 | Putative fumarate hydratase | Smlt3222 | 0.000 | <0.01 |
| B2FLG0 | Putative ferredoxin--NADP reductase | fpr | 0.000 | 0.24 |
| B2FLG2 | Putative peptidyl dipeptidase/oligopeptidase | Smlt3229 | 0.009 | 0.20 |
| B2FLG5 | Uncharacterized protein | Smlt3232 | 0.007 | 2.25 |
| B2FLG9 | Putative peptide transport protein | Smlt3236 | 0.033 | 2.07 |
| B2FLM4 | Biotin synthase | bioB | 0.000 | <0.01 |
| B2FLN3 | tRNA uridine 5-carboxymethylaminomethyl modification enzyme MnmG | mnmG | 0.000 | <0.01 |
| B2FLP0 | Putative transmembrane Na+/H+ antiport transporter | Smlt0555 | 0.016 | 5.04 |
| B2FLR0 | Putative 3-oxoacyl-[acyl-carrier-protein] synthase I | fabB | 0.014 | 0.43 |
| B2FLR1 | 3-hydroxydecanoyl-[acyl-carrier-protein] dehydratase | fabA | 0.004 | <0.01 |
| B2FLR6 | Uncharacterized protein | Smlt0583 | 0.001 | <0.01 |
| B2FLS9 | Histidine kinase | Smlt0596 | 0.000 | 12.16 |
| B2FLT0 | Putative two-component response regulator transcriptional regulatory protein | Smlt0597 | 0.002 | 2.23 |
| B2FLT5 | Putative TonB dependent receptor | Smlt0602 | 0.003 | 12.19 |
| B2FLT7 | Putative ATP-dependent RNA helicase 1 | rhlE1 | 0.000 | <0.01 |
| B2FLT8 | Putative fumarylacetoacetate (FAA) hydrolase family protein | Smlt0608 | 0.000 | <0.01 |
| B2FLU2 | Putative PilU protein (Twitching motility protein) | pilU | 0.001 | <0.01 |
| B2FLU9 | Putative ABC transporter component, polysaccharide related | wzt | 0.000 | 2.48 |
| B2FLV2 | Putative glycosyl transferase | wxocA | 0.001 | 2.84 |
| B2FLV9 | Putative FMN amine oxidoreductase | Smlt0629 | 0.001 | 2.83 |
| B2FLW0 | Putative transmembrane anchor NAD-dependent epimerase/dehydratase/dehydrogenase | Smlt0630 | 0.001 | 2.27 |
| B2FLW2 | Chaperone protein HtpG | htpG | 0.001 | 0.26 |
| B2FLW5 | Conserved hypothetical exported protein | Smlt1812 | 0.008 | 2.52 |
| B2FLW8 | Putative metallo-beta-lactamase superfamily protein | Smlt1815 | 0.000 | <0.01 |
| B2FLX2 | Uracil phosphoribosyltransferase | upp | 0.041 | 0.29 |
| B2FLX9 | Putative outer membrane lipoprotein | Smlt1826 | 0.002 | 4.48 |
| B2FLY0 | Putative LysR-family transcriptional regulator | Smlt1827 | 0.000 | <0.01 |
| B2FLZ7 | Putative modification methylase | Smlt1844 | 0.034 | <0.01 |
| B2FLZ8 | Putative modification methylase | Smlt1844A | 0.001 | <0.01 |
| B2FM02 | Uncharacterized protein | Smlt1846A | 0.000 | <0.01 |
| B2FM03 | Uncharacterized protein | Smlt1846B | 0.006 | 0.17 |
| B2FM53 | Uncharacterized protein | Smlt1897 | 0.000 | <0.01 |
| B2FM59 | Uncharacterized protein | Smlt1905 | 0.000 | <0.01 |
| B2FM60 | Putative phage-related protein | Smlt1906 | 0.002 | <0.01 |
| B2FM92 | Ribonuclease E | rnE | 0.005 | 0.31 |
| B2FM97 | Putative substrate-binding component of ABC transporter | Smlt3252 | 0.003 | 5.42 |
| B2FMA5 | Ribosomal large subunit pseudouridine synthase B | rluB | 0.029 | 0.33 |
| B2FMB8 | Putative cytochrome C-type biogenesis protein | dsbE | 0.009 | 2.56 |
| B2FMC0 | Putative cytochrome C-type biogenesis protein | cycH | 0.001 | 3.01 |
| B2FMC3 | Putative transmembrane ABC transporter ATP-binding protein, cytochrome related | cydC | 0.043 | 1.80 |
| B2FMC4 | Putative ABC transporter ATP-binding protein, cytochrome related | cydD | 0.000 | <0.01 |
| B2FMC5 | Putative cytochrome D ubiquinol oxidase subunit I | cydA | 0.008 | 0.33 |
| B2FMC6 | Putative transmembrane cytochrome bd-II oxidase subunit II | appB | 0.000 | <0.01 |
| B2FMD3 | Gamma-glutamyl phosphate reductase | proA | 0.002 | <0.01 |
| B2FMD6 | Argininosuccinate lyase | argH | 0.000 | <0.01 |
| B2FME5 | Cysteine--tRNA ligase | cysS | 0.000 | <0.01 |
| B2FMF1 | Putative exported peptidase | Smlt3311 | 0.015 | 3.54 |
| B2FMF2 | Uncharacterized protein | Smlt3312 | 0.000 | 2.81 |
| B2FMF3 | Phosphoglycolate phosphatase | Smlt3313 | 0.000 | <0.01 |
| B2FMG2 | Uncharacterized protein | Smlt4479 | 0.000 | <0.01 |
| B2FMI0 | Delta-aminolevulinic acid dehydratase | hemB | 0.013 | <0.01 |
| B2FMI1 | Putative transmembrane protein | Smlt4498 | 0.004 | 1.73 |
| B2FMI6 | Putative exported dipeptidyl peptidase IV | Smlt4503 | 0.006 | 0.16 |
| B2FMI7 | Putative glutathione S transferase | Smlt4504 | 0.000 | <0.01 |
| B2FMI8 | UPF0114 protein Smlt4505 | Smlt4505 | 0.037 | 2.80 |
| B2FMK0 | Putative oxidoreductase | Smlt4517 | 0.000 | <0.01 |
| B2FMK1 | Putative ABC transporter ATP-binding protein | Smlt4518 | 0.001 | 2.33 |
| B2FMK7 | Putative ATP-dependent helicase | hrpB | 0.015 | <0.01 |
| B2FML3 | Uncharacterized protein | Smlt4532 | 0.001 | <0.01 |
| B2FML4 | RNA-splicing ligase RtcB | Smlt4533 | 0.001 | <0.01 |
| B2FMM9 | Putative acyl-CoA synthetase | Smlt4550 | 0.000 | <0.01 |
| B2FMN2 | Uncharacterized protein | Smlt0631 | 0.001 | 2.89 |
| B2FMN3 | Putative transmembrane anchor short-chain dehydrogenase | Smlt0632 | 0.007 | 2.25 |
| B2FMN4 | Putative FAD-binding oxidoreductase | Smlt0633 | 0.005 | 2.84 |
| B2FMP0 | Uncharacterized protein | Smlt0639 | 0.000 | <0.01 |
| B2FMP7 | dTDP-glucose 4,6-dehydratase | rmlB | 0.012 | 0.34 |
| B2FMQ4 | Putative helicase | Smlt0654 | 0.007 | <0.01 |
| B2FMQ7 | Putative glycosyltransferase protein | wxocA | 0.002 | 1.96 |
| B2FMQ9 | Uncharacterized protein | Smlt0659 | 0.000 | 9.74 |
| B2FMR0 | Uncharacterized protein | Smlt0660 | 0.001 | 3.78 |
| B2FMR5 | Proline--tRNA ligase | proS | 0.000 | <0.01 |
| B2FMR8 | Putative transmembrane CDP-diacylglycerol--serine O-phosphatidyltransferase | Smlt0669 | 0.037 | 2.71 |
| B2FMS2 | Valine--tRNA ligase | valS | 0.003 | 0.29 |
| B2FMS5 | Putative transmembrane permease protein | Smlt0676 | 0.003 | 2.56 |
| B2FMS6 | Putative transmembrane permease protein | Smlt0677 | 0.000 | 2.91 |
| B2FMT0 | Putative thiol:disulfide interchange protein DsbC | dsbC | 0.011 | 2.57 |
| B2FMX8 | Outer membrane protein assembly factor BamE | bamE | 0.009 | 2.89 |
| B2FMY1 | Putative LysR family transcriptional regulator | Smlt1988 | 0.000 | <0.01 |
| B2FMY2 | DNA repair protein RecN | Smlt1989 | 0.004 | <0.01 |
| B2FMY3 | Heat-inducible transcription repressor HrcA | hrcA | 0.000 | <0.01 |
| B2FMY5 | Chaperone protein DnaK | dnaK | 0.008 | 1.62 |
| B2FMZ2 | Putative ABC transporter ATP-binding protein | Smlt2001 | 0.032 | 1.57 |
| B2FMZ9 | S-adenosylmethionine:tRNA ribosyltransferase-isomerase | queA | 0.010 | 0.27 |
| B2FN03 | Protein-export membrane protein SecF | secF | 0.034 | 2.16 |
| B2FN05 | Putative transmembrane protein | Smlt2016 | 0.002 | <0.01 |
| B2FN13 | Adenine phosphoribosyltransferase | apt | 0.001 | <0.01 |
| B2FN17 | Putative ABC transporter ATP-binding protein | uup | 0.001 | 0.19 |
| B2FN23 | Putative transmembrane TraB family protein | Smlt2036 | 0.000 | 2.94 |
| B2FN26 | Putative deiminase | Smlt2039 | 0.000 | <0.01 |
| B2FN29 | Putative inositol-1-monophosphatase | suhB | 0.005 | 0.30 |
| B2FN30 | Protease HtpX | htpX | 0.002 | 3.92 |
| B2FN33 | Putative poly-hydroxy-butyrate synthesis protein | Smlt3326 | 0.002 | 4.25 |
| B2FN37 | Putative N-acetylmuramoyl-L-alanine amidase | Smlt3330 | 0.004 | 2.04 |
| B2FN46 | Uncharacterized protein | Smlt3339 | 0.000 | <0.01 |
| B2FN47 | Putative TonB dependent receptor | Smlt3340 | 0.003 | 0.31 |
| B2FN54 | Conserved hypothetical exported protein | Smlt3351 | 0.003 | 5.26 |
| B2FN59 | Conserved hypothetical repetitive protein | Smlt3358 | 0.003 | 6.07 |
| B2FN75 | Phenylalanine--tRNA ligase alpha subunit | pheS | 0.016 | 0.34 |
| B2FN79 | Threonine--tRNA ligase | thrS | 0.039 | 0.45 |
| B2FN86 | Polyribonucleotide nucleotidyltransferase | pnp | 0.001 | 0.33 |
| B2FN90 | Translation initiation factor IF-2 | infB | 0.011 | 0.35 |
| B2FN91 | Putative N utilization substance protein A | nusA | 0.045 | 0.44 |
| B2FN97 | Putative transmembrane ankyrin repeat protein | Smlt4553 | 0.002 | 2.45 |
| B2FNA9 | Putative phenylalanine and histidine ammonia-lyase | Smlt4565 | 0.000 | <0.01 |
| B2FNB2 | Putative AMP-binding enzyme | Smlt4568 | 0.000 | <0.01 |
| B2FNC5 | Putative dipeptidyl peptidase | Smlt4581 | 0.013 | 2.10 |
| B2FND3 | Putative DNA polymerase I | polA | 0.003 | 0.22 |
| B2FND7 | Uncharacterized protein | Smlt4593 | 0.038 | 0.55 |
| B2FND9 | DNA helicase | uvrD | 0.007 | 0.05 |
| B2FNE5 | Putative exonuclease | Smlt4602 | 0.000 | <0.01 |
| B2FNF2 | Uncharacterized protein | Smlt4609 | 0.000 | <0.01 |
| B2FNF8 | Putative chromosome partitioning protein | parA | 0.009 | 0.34 |
| B2FNG2 | Putative endonuclease/exonuclease/phosphatase family protein | Smlt4619 | 0.001 | <0.01 |
| B2FNH0 | Conserved hypothetical exported protein | Smlt4628 | 0.000 | 0.51 |
| B2FNH1 | Putative chaperone heat shock Hsp70 protein | hscC | 0.000 | <0.01 |
| B2FNI3 | Putative general secretion pathway protein D | xpsD | 0.009 | 4.39 |
| B2FNJ0 | Putative fimbrial adhesin protein | smf-1 | 0.006 | 5.88 |
| B2FNJ2 | Putative outer membrane usher protein mrkc | mrkC | 0.011 | 6.08 |
| B2FNJ3 | Putative fimbria adhesin protein | Smlt0709 | 0.000 | 5.24 |
| B2FNK0 | Putative ABC transporter component protein | Smlt0716 | 0.000 | 0.23 |
| B2FNL0 | Uncharacterized protein | Smlt0726 | 0.003 | 0.47 |
| B2FNM5 | Putative LppC family lipoprotein | Smlt0742 | 0.007 | 3.46 |
| B2FNN2 | Putative penicillin-binding protein 3 | ftsI | 0.004 | 2.54 |
| B2FNN4 | UDP-N-acetylmuramoyl-tripeptide--D-alanyl-D-alanine ligase | murF | 0.009 | 0.21 |
| B2FNN7 | UDP-N-acetylglucosamine--N-acetylmuramyl-(pentapeptide) pyrophosphoryl-undecaprenol N-acetylglucosamine transferase | murG | 0.003 | 2.68 |
| B2FNP0 | Cell division protein FtsQ | ftsQ | 0.003 | 3.34 |
| B2FNP8 | Uncharacterized protein | Smlt2046 | 0.000 | 2.55 |
| B2FNQ0 | Putative epimerase/dehydratase polysaccharide-related biosynthesis protein | wbiI | 0.008 | 1.69 |
| B2FNQ5 | Nucleoside diphosphate kinase | ndk | 0.001 | 0.28 |
| B2FNQ9 | Putative transmembrane protein | Smlt2058 | 0.000 | 5.90 |
| B2FNR0 | Outer membrane protein assembly factor BamB | bamB | 0.004 | 3.16 |
| B2FNR9 | Bifunctional protein FolD | folD | 0.000 | <0.01 |
| B2FNS0 | Inosine-5'-monophosphate dehydrogenase | guaB | 0.016 | 0.51 |
| B2FNS1 | GMP synthase [glutamine-hydrolyzing] | guaA | 0.032 | 0.28 |
| B2FNS5 | Uncharacterized protein | Smlt2077A | 0.041 | <0.01 |
| B2FNX3 | Putative NADH-ubiquinone oxidoreductase, 75 kDa subunit | nuoG | 0.005 | 1.84 |
| B2FNY0 | Putative general secretory pathway protein-export membrane protein | secG | 0.005 | 2.18 |
| B2FNY2 | Putative flavonol synthase/dioxygenase | Smlt3409 | 0.002 | 0.21 |
| B2FNY4 | Putative lipopolysaccharide core biosynthesis glycosyl transferase | Smlt3411 | 0.003 | 2.15 |
| B2FNY7 | Phosphoglucosamine mutase | glmM | 0.000 | <0.01 |
| B2FNY8 | Acetyl-coenzyme A carboxylase carboxyl transferase subunit beta | accD | 0.015 | 0.64 |
| B2FNZ1 | Tryptophan synthase beta chain | trpB | 0.023 | 0.38 |
| B2FNZ8 | Putative fimV protein | fimV | 0.004 | 3.14 |
| B2FP03 | Putative exported Sco1/SenC family protein | Smlt3431 | 0.001 | 3.14 |
| B2FP04 | Phosphatidylserine decarboxylase proenzyme | psd | 0.008 | 3.05 |
| B2FP06 | Putative membrane-bound lytic murein transglycosylase d | mltD | 0.002 | 3.49 |
| B2FP10 | Ribosomal protein S12 methylthiotransferase RimO | rimO | 0.000 | <0.01 |
| B2FP15 | Putative ATP-binding protein | Smlt3443 | 0.001 | <0.01 |
| B2FP17 | Putative TonB dependent receptor protein | Smlt3446 | 0.029 | 1.92 |
| B2FP18 | Putative endopeptidase O | pepO | 0.002 | 2.45 |
| B2FP19 | Putative TonB dependent receptor protein | Smlt3449 | 0.047 | 0.44 |
| B2FP20 | Putative exported endopeptidase | Smlt3450 | 0.015 | 2.27 |
| B2FP30 | Putative lipoprotein | Smlt3460 | 0.003 | 4.37 |
| B2FP33 | Putative alkaline phosphatase | Smlt3463 | 0.012 | 2.49 |
| B2FP46 | Putative surface antigen exported protein | Smlt4633 | 0.004 | 2.70 |
| B2FP49 | Putative type II/IV secretion system protein | Smlt4636 | 0.007 | <0.01 |
| B2FP53 | Sec-independent protein translocase protein TatB | tatB | 0.000 | 5.94 |
| B2FP62 | Putative transglycosylase protein | Smlt4650 | 0.004 | 3.61 |
| B2FP71 | GTP cyclohydrolase 1 | folE | 0.000 | <0.01 |
| B2FP82 | Putative ABC transporter ATP-binding protein | Smlt4670 | 0.000 | 2.55 |
| B2FP84 | Putative transmembrane mce related protein | Smlt4672 | 0.001 | 3.80 |
| B2FP85 | Putative ABC transporter toluene tolerance exported protein | Smlt4673 | 0.007 | 2.41 |
| B2FP87 | Putative intercellular spreading VacJ lipoprotein | Smlt4675 | 0.003 | 3.95 |
| B2FP90 | Putative RmuC family protein | Smlt4678 | 0.001 | 1.77 |
| B2FP98 | Putative dipeptidase | Smlt4686 | 0.029 | <0.01 |
| B2FPA5 | Membrane protein insertase YidC | yidC | 0.004 | 1.81 |
| B2FPA8 | Cell division protein FtsZ | ftsZ | 0.011 | 0.42 |
| B2FPA9 | UDP-3-O-[3-hydroxymyristoyl] N-acetylglucosamine deacetylase | lpxC | 0.002 | 0.11 |
| B2FPB2 | Protein translocase subunit SecA | secA | 0.008 | 2.19 |
| B2FPC7 | S-adenosylmethionine synthase | metK | 0.000 | 2.80 |
| B2FPE0 | Putative aldo/keto reductase family protein | Smlt0793 | 0.000 | <0.01 |
| B2FPE2 | Putative exported heme receptor protein | huvA | 0.002 | 79.94 |
| B2FPF6 | Putative LemA family protein | lemA | 0.000 | 11.08 |
| B2FPF9 | Prolipoprotein diacylglyceryl transferase | lgt | 0.002 | 2.57 |
| B2FPG7 | Chaperone SurA | surA | 0.001 | 6.68 |
| B2FPG8 | LPS-assembly protein LptD | lptD | 0.014 | 2.53 |
| B2FPH2 | Putative 2-octaprenyl-6-methoxyphenol hydroxylase | visB | 0.000 | <0.01 |
| B2FPI7 | Dihydroorotate dehydrogenase (quinone) | pyrD | 0.000 | 2.45 |
| B2FPJ1 | Putative aldehyde dehydrogenase | Smlt2132 | 0.009 | 0.20 |
| B2FPJ6 | Putative universal stress family protein | Smlt2137 | 0.007 | 2.78 |
| B2FPK0 | Putative repetitive protein with two-component sensor and regulator motifs | Smlt2141 | 0.000 | <0.01 |
| B2FPK3 | Putative two component system histidine kinase/response regulator fusion protein | Smlt2144 | 0.020 | <0.01 |
| B2FPN2 | Putative copper-transporting P-type ATPase | actP | 0.036 | 2.01 |
| B2FPN5 | Putative TonB-dependent outer membrane receptor protein | Smlt2179 | 0.039 | 0.50 |
| B2FPQ0 | Putative D-3-phosphoglycerate dehydrogenase | serA | 0.014 | 0.21 |
| B2FPQ1 | Putative FAD binding oxidoreductase | Smlt2196 | 0.000 | <0.01 |
| B2FPQ6 | Putative ribonuclease G | rnG | 0.033 | <0.01 |
| B2FPR6 | LPS-assembly lipoprotein LptE | lptE | 0.000 | 4.31 |
| B2FPR7 | Leucine--tRNA ligase | leuS | 0.000 | <0.01 |
| B2FPR9 | Putative thioredoxin protein | Smlt3487 | 0.001 | 0.10 |
| B2FPS4 | Putative transmembrane protein | Smlt3492 | 0.025 | 1.90 |
| B2FPS6 | Uncharacterized protein | Smlt3494 | 0.020 | 1.96 |
| B2FPT0 | Methionine--tRNA ligase | metG | 0.010 | 0.61 |
| B2FPU1 | Putative heat shock chaperone protein | Smlt3509 | 0.000 | <0.01 |
| B2FPV0 | Uncharacterized protein | Smlt3519 | 0.000 | <0.01 |
| B2FPV6 | Putative thiolase | Smlt3525 | 0.005 | <0.01 |
| B2FPW8 | Putative hypoxanthine phosphoribosyltransferase | hpt | 0.003 | <0.01 |
| B2FPY1 | Putative signal peptidase I | lepB | 0.010 | 1.77 |
| B2FPZ4 | Putative exodeoxyribonuclease | Smlt0848 | 0.000 | <0.01 |
| B2FQ05 | Branched-chain-amino-acid aminotransferase | ilvE | 0.029 | 0.40 |
| B2FQ07 | Putative asparaginase | Smlt0862 | 0.027 | 0.40 |
| B2FQ10 | UPF0761 membrane protein Smlt0865 | Smlt0865 | 0.015 | 1.86 |
| B2FQ11 | Putative thioredoxin electron transport related protein | Smlt0866 | 0.001 | 3.65 |
| B2FQ15 | Glutamyl-tRNA reductase | hemA | 0.001 | <0.01 |
| B2FQ17 | Outer-membrane lipoprotein LolB | lolB | 0.001 | 5.51 |
| B2FQ19 | Ribose-phosphate pyrophosphokinase | prs | 0.031 | 0.24 |
| B2FQ22 | Ribosome-binding ATPase YchF | ychF | 0.004 | 0.16 |
| B2FQ32 | Protein translocase subunit SecE | secE | 0.010 | 1.87 |
| B2FQ38 | DNA-directed RNA polymerase subunit beta | rpoB | 0.011 | 0.32 |
| B2FQ39 | DNA-directed RNA polymerase subunit beta' | rpoC | 0.012 | 0.32 |
| B2FQ42 | Elongation factor G | fusA | 0.040 | 0.51 |
| B2FQ57 | Uncharacterized protein | Smlt2204 | 0.002 | 6.14 |
| B2FQ59 | Putative short-chain dehydrogenase family protein | Smlt2206 | 0.000 | <0.01 |
| B2FQ84 | Lysine--tRNA ligase | lysS | 0.001 | <0.01 |
| B2FQ85 | Putative two-component response regulator transcriptional regulatory protein, regulator of pathogenicity factors | rpfG | 0.000 | <0.01 |
| B2FQ88 | Putative long-chain-fatty-acid--CoA ligase | fadD | 0.007 | 2.07 |
| B2FQ89 | Oxygen-dependent choline dehydrogenase | betA | 0.001 | 4.96 |
| B2FQ93 | Aconitate hydratase | acn | 0.003 | <0.01 |
| B2FQ97 | Aconitate hydratase 2 | acnB | 0.000 | 0.16 |
| B2FQC4 | Elongation factor 4 | lepA | 0.003 | 0.30 |
| B2FQC5 | Putative subfamily S1C unassigned peptidase | Smlt3553 | 0.000 | 3.95 |
| B2FQC8 | Putative enoyl-CoA hydratasee | Smlt3556 | 0.001 | 4.39 |
| B2FQD3 | Putative carbon starvation protein A | cstA | 0.046 | 3.60 |
| B2FQE3 | Putative angiotensin-converting enzyme like peptidyl dipeptidase protein | Smlt3574 | 0.022 | 1.61 |
| B2FQE7 | Glycine dehydrogenase (decarboxylating) | gcvP | 0.000 | <0.01 |
| B2FQG0 | Adenylosuccinate synthetase | purA | 0.008 | <0.01 |
| B2FQG3 | Protein HflC | Smlt3595 | 0.001 | 2.97 |
| B2FQG4 | Putative HflK protein | hflK | 0.002 | 2.48 |
| B2FQG7 | Putative curved DNA-binding protein | cbpA | 0.013 | 0.28 |
| B2FQH7 | Citrate synthase | prpC | 0.000 | <0.01 |
| B2FQH9 | Putative sigma 54 binding transcriptional regulatory protein | Smlt3611 | 0.000 | <0.01 |
| B2FQI7 | Shikimate kinase | aroK | 0.000 | <0.01 |
| B2FQK4 | Protein translocase subunit SecY | secY | 0.043 | 1.69 |
| B2FQK8 | DNA-directed RNA polymerase subunit alpha | rpoA | 0.006 | 0.39 |
| B2FQL6 | Putative GTP-binding protein | typA | 0.000 | 0.20 |
| B2FQL7 | Peptidyl-prolyl cis-trans isomerase | ppi | 0.043 | 2.35 |
| B2FQL8 | Malate dehydrogenase | mdh | 0.004 | 2.67 |
| B2FQM8 | Putative outer membrane protein | Smlt0955 | 0.012 | 2.18 |
| B2FQN2 | Putative 2-amino-3-ketobutyrate coenzyme A ligase | kbl | 0.000 | <0.01 |
| B2FQN3 | Uncharacterized protein | Smlt0960 | 0.002 | 0.06 |
| B2FQN7 | Putative FolC bifunctional protein [includes: folylpolyglutamate synthase and dihydrofolate synthase] | folC | 0.010 | 0.30 |
| B2FQN8 | Conserved hypothetical exported protein | Smlt0965 | 0.000 | 3.82 |
| B2FQP8 | Putative exopolyphosphatase | ppx | 0.005 | 3.72 |
| B2FQQ0 | Histidine kinase | phoR | 0.001 | 5.52 |
| B2FQQ2 | Putative exported peptidase | Smlt0979 | 0.020 | 4.13 |
| B2FQQ5 | Putative isocitrate/isopropylmalate dehydrogenase | Smlt0982 | 0.022 | 0.40 |
| B2FQR3 | ATP-dependent Clp protease ATP-binding subunit ClpX | clpX | 0.035 | 0.51 |
| B2FQR4 | Lon protease | lon | 0.048 | 0.55 |
| B2FQS5 | Putative flagellar basal body-associated protein FliL | fliL | 0.002 | 7.32 |
| B2FQT7 | Putative nitrogen regulation protein NR(I) | Smlt2295 | 0.000 | <0.01 |
| B2FQU5 | Putative flagellin | fliC | 0.000 | 19.57 |
| B2FQU6 | Putative flagellin | flaA | 0.000 | 20.46 |
| B2FQU7 | Putative motility flagellin protein | Smlt2306 | 0.000 | 18.49 |
| B2FQW3 | Histidine kinase | Smlt2322 | 0.003 | <0.01 |
| B2FQW4 | Putative GGDEF EAL response regulator signalling protein | Smlt2323 | 0.000 | <0.01 |
| B2FQX1 | High frequency lysogenization protein HflD homolog | hflD | 0.000 | 3.39 |
| B2FQX7 | Putative ATP-dependent Clp protease ATP-binding subunit | clpA | 0.036 | 0.55 |
| B2FQY4 | Putative cell division protein | ftsK | 0.003 | 2.20 |
| B2FQY7 | Putative ATPase | Smlt2348 | 0.001 | <0.01 |
| B2FQZ4 | Putative L-serine dehydratase I | sdaA | 0.000 | <0.01 |
| B2FR00 | Peptide chain release factor 3 | prfC | 0.000 | 0.09 |
| B2FR06 | 33 kDa chaperonin | Smlt3643 | 0.000 | <0.01 |
| B2FR11 | Putative acyl-coenzyme A dehydrogenase | fadE | 0.004 | 3.06 |
| B2FR23 | Putative transmembrane protein | Smlt3660 | 0.002 | 2.11 |
| B2FR33 | Putative gliding motility sensor histidine kinase response regulator fusion transcriptional regulatory protein | pilL | 0.000 | <0.01 |
| B2FR34 | Putative pilus biogenesis protein PilJ/methyl accepting chemotaxis protein | pilJ | 0.003 | 0.37 |
| B2FR35 | Putative pilus biogenesis protein | pilI | 0.016 | <0.01 |
| B2FR41 | Putative ATP-dependent DNA helicase-related protein | Smlt3678 | 0.002 | 0.08 |
| B2FR42 | Conserved hypothetical exported protein | Smlt3680 | 0.003 | 5.70 |
| B2FR43 | Putative penicillin-binding protein 1B | mrcB | 0.001 | 4.89 |
| B2FR44 | Putative glycosyl transferase | Smlt3682 | 0.044 | 2.33 |
| B2FR45 | Putative O-antigen related protein | wbpV | 0.001 | 2.61 |
| B2FR47 | Putative GTP pyrophosphokinase | relA | 0.022 | 0.23 |
| B2FR49 | Putative ATP-dependent RNA helicase HrpA | hrpA | 0.000 | 0.04 |
| B2FR51 | Putative ATP-dependent DNA helicase | recQ | 0.005 | <0.01 |
| B2FR55 | Peptidylprolyl isomerase | Smlt0993 | 0.003 | 4.40 |
| B2FR61 | Putative phosphatase | Smlt0999 | 0.001 | 6.22 |
| B2FR62 | Putative autotransporter | Smlt1001 | 0.002 | <0.01 |
| B2FR64 | Putative ADP-heptose--LPS heptosyltransferase II | waaF | 0.006 | 3.08 |
| B2FR73 | Putative restriction-modification system methyltransferase | Smlt1013 | 0.000 | <0.01 |
| B2FR74 | Putative DNA polymerase III subunit Tau | dnaX | 0.011 | 0.30 |
| B2FR78 | Putative outer membrane lipoprotein | Smlt1018 | 0.004 | 5.01 |
| B2FR87 | Uncharacterized protein | Smlt1027 | 0.012 | 2.21 |
| B2FR91 | 3-oxoacyl-[acyl-carrier-protein] synthase 2 | fabF | 0.001 | <0.01 |
| B2FR92 | Putative chorismate binding protein | Smlt1032 | 0.000 | <0.01 |
| B2FR93 | Putative aminodeoxychorismate lyase | Smlt1034 | 0.000 | 4.24 |
| B2FRA4 | Putative bacteriophage major tail sheath protein | Smlt1045 | 0.001 | <0.01 |
| B2FRC4 | Putative TonB-dependent receptor | Smlt1067 | 0.006 | 33.70 |
| B2FRD8 | Conserved hypothetical exported protein | Smlt1082 | 0.027 | 5.42 |
| B2FRE5 | Putative binding-protein-dependent transport lipoprotein | Smlt2355 | 0.000 | 169.00 |
| B2FRM8 | Putative peptidoglycan-associated lipoprotein | Smlt3703 | 0.008 | 2.73 |
| B2FRM9 | Protein TolB | tolB | 0.003 | 2.98 |
| B2FRN0 | Putative TolA transmembrane protein | tolA | 0.000 | 4.97 |
| B2FRN1 | Putative TolR-related protein | tolR | 0.001 | 4.41 |
| B2FRN2 | Putative TolQ transport transmembrane protein | tolQ | 0.003 | 2.74 |
| B2FRN4 | Holliday junction ATP-dependent DNA helicase RuvB | ruvB | 0.001 | <0.01 |
| B2FRN5 | Probable potassium transport system protein kup | kup | 0.021 | 1.56 |
| B2FRN6 | Holliday junction ATP-dependent DNA helicase RuvA | ruvA | 0.001 | <0.01 |
| B2FRN7 | Crossover junction endodeoxyribonuclease RuvC | ruvC | 0.000 | <0.01 |
| B2FRP3 | Uncharacterized protein | Smlt3720 | 0.015 | 4.55 |
| B2FRP7 | Putative ABC transport system, ATP-binding protein | Smlt3724 | 0.002 | 0.21 |
| B2FRP8 | Putative TonB dependent receptor | Smlt3725 | 0.007 | 5.51 |
| B2FRP9 | Putative transmembrane transport protein | Smlt3726 | 0.003 | 2.50 |
| B2FRQ4 | Putative heat shock chaperone ClpB | clpB | 0.003 | 0.34 |
| B2FRR0 | Putative lipoprotein | Smlt3739 | 0.004 | 9.55 |
| B2FRR1 | Putative TonB dependent receptor | Smlt3740 | 0.007 | 9.13 |
| B2FRR9 | Outer membrane protein assembly factor BamD | comL | 0.012 | 3.49 |
| B2FRS1 | Putative glutamine-dependent NAD synthase | nadE | 0.000 | <0.01 |
| B2FRS5 | Putative transcriptional regulatory protein | hydG | 0.002 | 0.25 |
| B2FRS6 | Putative type IV pilus assembly protein | pilF | 0.005 | 0.28 |
| B2FRS9 | Putative pilus-assembly protein | pilG | 0.001 | <0.01 |
| B2FRT4 | Histidine kinase | Smlt3765 | 0.003 | 3.56 |
| B2FRU2 | Putative twitching mobility protein | pilT | 0.000 | <0.01 |
| B2FRU3 | Putative pilus retraction protein | pilU | 0.000 | <0.01 |
| B2FRW5 | Putative ABC transporter, ATP-binding protein | Smlt1113 | 0.005 | 2.57 |
| B2FRW6 | Putative OstA family protein | Smlt1114 | 0.002 | 7.96 |
| B2FRX1 | UDP-N-acetylglucosamine 1-carboxyvinyltransferase | murA | 0.002 | <0.01 |
| B2FRX8 | Phosphoribosylformylglycinamidine cyclo-ligase | purM | 0.014 | 0.35 |
| B2FRX9 | Conserved hypothetical exported protein | Smlt1127 | 0.002 | 3.16 |
| B2FRY0 | Putative transmembrane protein | Smlt1128 | 0.012 | 2.85 |
| B2FRY5 | Putative peptidase | Smlt1133 | 0.000 | <0.01 |
| B2FRZ0 | Putative oxidoreductase | Smlt1138 | 0.000 | 5.17 |
| B2FRZ4 | Putative transmembrane PepSY family protein | Smlt1142 | 0.000 | 4.45 |
| B2FRZ5 | Putative exogenous ferric siderophore receptor | bfrA | 0.016 | 4.65 |
| B2FRZ9 | Putative iron transport receptor protein | Smlt1148 | 0.005 | 5.67 |
| B2FS04 | Cysteine desulfurase | Smlt1153 | 0.006 | <0.01 |
| B2FS15 | Putative histone-like protein | Smlt1164 | 0.005 | 2.27 |
| B2FS54 | Putative integrase | Smlt2472 | 0.000 | <0.01 |
| B2FS56 | Uncharacterized protein | Smlt2475 | 0.000 | <0.01 |
| B2FSC8 | Putative outer membrane esterase | Smlt3773 | 0.006 | 3.19 |
| B2FSD1 | tRNA (guanine-N(7)-)-methyltransferase | trmB | 0.001 | <0.01 |
| B2FSD7 | Large-conductance mechanosensitive channel | mscL | 0.005 | 2.68 |
| B2FSE3 | Putative HlyD-family secretion protein | smeM | 0.011 | 1.71 |
| B2FSE4 | Putative TonB dependent receptor | Smlt3789 | 0.004 | 28.72 |
| B2FSF1 | Pyruvate kinase | pykA | 0.003 | 0.22 |
| B2FSF4 | Putative transmembrane protein | Smlt3802 | 0.003 | 1.89 |
| B2FSF6 | Glyceraldehyde-3-phosphate dehydrogenase | gap | 0.003 | 4.36 |
| B2FSF7 | Putative outer membrane Omp family protein | Smlt3805 | 0.049 | 0.46 |
| B2FSF8 | Putative endonuclease P1 | Smlt3806 | 0.028 | 3.59 |
| B2FSG4 | Uncharacterized protein | Smlt3812 | 0.010 | 2.73 |
| B2FSG5 | Transketolase | tktA | 0.004 | 0.16 |
| B2FSG6 | Putative transmembrane sodium-dicarboxylate family transporter protein | Smlt3814 | 0.024 | 2.24 |
| B2FSG9 | Putative von Willebrand factor-like protein | Smlt3817 | 0.002 | 3.38 |
| B2FSH1 | Uncharacterized protein | Smlt3819 | 0.002 | 3.61 |
| B2FSH3 | Putative type II/III pilus secretin | pilQ | 0.003 | <0.01 |
| B2FSH4 | Putative pilP protein (Type 4 fimbrial biogenesis protein) | pilP | 0.001 | <0.01 |
| B2FSH5 | Putative PilO protein (Type 4 fimbrial biogenesis protein PilO) | pilO | 0.003 | <0.01 |
| B2FSH6 | Putative PilN protein (Type 4 fimbrial biogenesis protein) | pilN | 0.007 | <0.01 |
| B2FSH7 | Putative PilM protein (Type 4 fimbrial biogenesis protein) | pilM | 0.002 | <0.01 |
| B2FSH8 | Putative penicillin-binding protein | mrcA | 0.001 | 2.74 |
| B2FSI7 | Citrate synthase | gltA | 0.014 | 2.10 |
| B2FSI9 | Putative nucleoside hydrolase | Smlt3837 | 0.000 | <0.01 |
| B2FSJ3 | Uncharacterized protein | Smlt1169 | 0.000 | <0.01 |
| B2FSJ9 | Putative TonB dependent receptor protein | Smlt1175 | 0.005 | <0.01 |
| B2FSL1 | Single-stranded DNA-binding protein | ssb | 0.033 | 0.38 |
| B2FSP6 | Putative inosine-uridine preferring nucleoside hydrolase | Smlt1222 | 0.001 | <0.01 |
| B2FSQ7 | Putative TonB-dependent receptor for Fe(III)-coprogen, Fe(III)-ferrioxamine B and Fe(III)-rhodotrulic acid | Smlt1233 | 0.005 | 47.32 |
| B2FSQ8 | Probable malate:quinone oxidoreductase | mqo | 0.009 | 2.64 |
| B2FSS4 | Putative multiphosphoryl transfer protein | Smlt2556 | 0.040 | 4.53 |
| B2FSS5 | Phosphofructokinase | fruK | 0.000 | <0.01 |
| B2FSS6 | Putative PTS system, fructose-specific IIBC component | fruA | 0.018 | 3.11 |
| B2FSS7 | Putative outer membrane regulator of pathogenicity factors protein | rpfN | 0.002 | 3.35 |
| B2FST2 | Putative TonB dependent receptor | Smlt2566 | 0.026 | 2.98 |
| B2FT04 | Putative guanosine-3',5'-bis(Diphosphate) 3'-pyrophosphohydrolase | spoT | 0.014 | 0.22 |
| B2FT06 | Guanylate kinase | gmk | 0.000 | <0.01 |
| B2FT17 | Ribonuclease PH | rph | 0.030 | <0.01 |
| B2FT30 | Ribose-5-phosphate isomerase A | rpiA | 0.050 | <0.01 |
| B2FT31 | Putative ACR family protein | Smlt3869 | 0.005 | 2.78 |
| B2FT35 | Glutamate-1-semialdehyde 2,1-aminomutase | hemL | 0.027 | 0.36 |
| B2FT36 | Putative azurin | Smlt3873A | 0.035 | 0.25 |
| B2FT45 | Putative transmembrane DedA family protein | Smlt3883 | 0.005 | 5.34 |
| B2FT63 | Inorganic pyrophosphatase | ppa | 0.049 | 2.60 |
| B2FT64 | Putative two component response regulator transcriptional regulatory protein | Smlt3903 | 0.000 | <0.01 |
| B2FT68 | Putative transcriptional regulator | Smlt3907 | 0.024 | 1.70 |
| B2FT83 | DNA gyrase subunit B | gyrB | 0.013 | 0.35 |
| B2FT85 | Putative peptidase | Smlt0007 | 0.003 | 3.95 |
| B2FT87 | Putative proline-rich TonB dependent receptor protein | Smlt0009 | 0.005 | 0.56 |
| B2FT88 | Putative biopolymer transport exbB protein | exbB1 | 0.001 | 2.91 |
| B2FT89 | Putative biopolymer transport ExbD1 protein | exbD1 | 0.001 | 3.05 |
| B2FT90 | Putative biopolymer transport ExbD2 protein | exbD2 | 0.007 | 3.43 |
| B2FT92 | Uncharacterized protein | Smlt0014 | 0.001 | 5.61 |
| B2FTA2 | Putative oligopeptide transporter | Smlt1245 | 0.012 | 2.22 |
| B2FTA3 | Putative exported peptidase | Smlt1246 | 0.003 | 4.59 |
| B2FTA6 | Cell division topological specificity factor | minE | 0.007 | 3.04 |
| B2FTA9 | Uncharacterized protein | Smlt1253 | 0.010 | 0.33 |
| B2FTB2 | Conserved hypothetical exported protein | Smlt1256 | 0.003 | 2.94 |
| B2FTB8 | Putative transmembrane protein | Smlt1263 | 0.039 | 3.37 |
| B2FTD0 | UvrABC system protein A | uvrA | 0.001 | <0.01 |
| B2FTD3 | GTPase Obg | obg | 0.005 | 0.25 |
| B2FTI8 | Putative lipid II flippase MurJ | mviN | 0.002 | 2.29 |
| B2FTI9 | Riboflavin biosynthesis protein | ribF | 0.004 | <0.01 |
| B2FTK0 | Putative two-component regulatory system family, response regulator protein | Smlt2645 | 0.011 | 3.44 |
| B2FTK7 | Proline iminopeptidase | pip | 0.000 | <0.01 |
| B2FTM0 | Putative TonB-dependent ferric siderophore receptor | Smlt2666 | 0.037 | 1.88 |
| B2FTS2 | Putative RND/Acr family transmembrane transporter | smeO | 0.001 | 5.22 |
| B2FTS7 | Putative 3-deoxy-D-manno-octulosonic-acid transferase | kdtA | 0.002 | 1.63 |
| B2FTS8 | Putative lipid A biosynthesis lauroyl acyltransferase | htrB | 0.007 | 2.18 |
| B2FTT5 | Putative NADP-dependent malic enzyme | maeB | 0.030 | 0.61 |
| B2FTU3 | Putative phosphate selective porin | Smlt3950 | 0.020 | 1.94 |
| B2FTU4 | Citrate transporter | citM | 0.016 | 1.80 |
| B2FTU5 | Putative acetoacetyl-CoA reductase | phbB | 0.009 | 2.89 |
| B2FTV8 | Conserved hypothetical exported protein | Smlt3965 | 0.001 | <0.01 |
| B2FTW0 | Putative DegV family transcriptional regulatory protein | Smlt3967 | 0.007 | 0.14 |
| B2FTY7 | Thiol:disulfide interchange protein | dsbA | 0.001 | 4.45 |
| B2FTY8 | Putative thiol:disulfide interchange protein | Smlt3994 | 0.002 | 3.71 |
| B2FTY9 | Putative endonuclease/exonuclease/phosphatase family protein | Smlt3995 | 0.002 | 0.25 |
| B2FU40 | Putative patatin-like phospholipase | Smlt0080 | 0.006 | 2.84 |
| B2FU51 | Thymidine kinase | tdk | 0.011 | <0.01 |
| B2FU54 | DNA helicase | rep | 0.000 | <0.01 |
| B2FU57 | Putative lipoprotein E (Outer membrane protein p4) | hel | 0.006 | 2.57 |
| B2FU60 | Glycerol-3-phosphate acyltransferase | plsB | 0.003 | 2.09 |
| B2FU61 | tRNA 2-thiocytidine biosynthesis protein TtcA | ttcA | 0.000 | <0.01 |
| B2FU78 | Putative helicase | lhr | 0.000 | <0.01 |
| B2FU86 | Uncharacterized protein | Smlt1345 | 0.041 | 1.83 |
| B2FU91 | Putative outer membrane autotransporter | Smlt1350 | 0.000 | <0.01 |
| B2FUA0 | Putative transmembrane ubiquinol oxidase subunit 2 | qoxA | 0.033 | 1.68 |
| B2FUA1 | Putative quinol oxidase subunit 1 | qoxB | 0.015 | 1.98 |
| B2FUA5 | DNA repair protein radA | radA | 0.000 | <0.01 |
| B2FUA6 | Putative chaperone protein HtpG (Heat shock protein HtpG) | htpG | 0.047 | 0.44 |
| B2FUA7 | Uncharacterized protein | Smlt1368 | 0.000 | <0.01 |
| B2FUB0 | Signal recognition particle protein | ffh | 0.020 | 0.47 |
| B2FUD9 | Putative HlyD family secretion protein | Smlt1406 | 0.002 | 3.56 |
| B2FUE8 | Putative diaminobutyrate--2-oxoglutarate aminotransferase | dat | 0.001 | 12.04 |
| B2FUK5 | Putative fumarate and nitrate reduction transcriptional regulatory protein | fnr2 | 0.000 | <0.01 |
| B2FUK6 | Putative oxygen-independent coproporphyrinogen III oxidase | Smlt2768 | 0.041 | <0.01 |
| B2FUL1 | Putative respiratory nitrate reductase subunit | narH | 0.000 | <0.01 |
| B2FUL2 | Putative respiratory nitrate reductase alpha chain | narG | 0.000 | 0.03 |
| B2FUN9 | Putative TonB-dependent receptor for Fe(III)-coprogen, Fe(III)-ferrioxamine B and Fe(III)-rhodotrulic acid | fhuE | 0.001 | 23.49 |
| B2FUP4 | Putative transglycosylase | Smlt4007 | 0.005 | 2.27 |
| B2FUQ7 | Putative phosphosugar-binding protein | Smlt4021 | 0.041 | 0.28 |
| B2FUR4 | Uncharacterized protein | Smlt4029 | 0.000 | <0.01 |
| B2FUS1 | Histidine kinase | Smlt4039 | 0.000 | 3.25 |
| B2FUS9 | Octanoyltransferase | lipB | 0.000 | <0.01 |
| B2FUT1 | Conserved hypothetical exported protein | Smlt4049 | 0.011 | 2.18 |
| B2FUT2 | Putative penicillin-binding protein | dacC | 0.001 | 4.12 |
| B2FUT3 | Putative exported rare lipoprotein A | rlpA | 0.007 | 4.18 |
| B2FUT4 | Putative murein hydrolase | mltB | 0.000 | 3.03 |
| B2FUT9 | Putative rod shape-determining protein | mreC | 0.000 | 3.01 |
| B2FUU4 | Uncharacterized protein | Smlt4063 | 0.003 | 4.39 |
| B2FUU5 | Putative PEPTIDASE | Smlt4064 | 0.003 | 3.79 |
| B2FUV1 | Putative multidrug resistance outer membrane protein | smeF | 0.010 | 2.85 |
| B2FUV3 | Putative acriflavin resistance protein A | smeD | 0.000 | 2.98 |
| B2FUV6 | ATP-dependent protease ATPase subunit HslU | hslU | 0.014 | 0.52 |
| B2FUW1 | Chromosomal replication initiator protein DnaA | dnaA | 0.002 | 0.24 |

**Table S3 Normalised proteomics data for mutant KLTV relative to K279a parent**

Abundance changes significantly according to a T-Test (P<0.05). Proteins that are >1.5 fold up and down regulated are highlighted green or red, respectively. Proteins are ordered based on Uniprot Accession number. Consecutive numbers are suggestive of operons. Where proteins are not detectable in one sample, it is not possible to give an accurate fold change so >100 or <0.01 are used to represent the actual change, which must be considerable.

| Accession | Description |  | T-Test _K279a vs KLTV_ | Fold _K279a vs KLTV_ |
| --- | --- | --- | --- | --- |
| B2FHA2 | Putative ferredoxin oxidoreductase | Smlt0136 | 0.039 | 0.13 |
| B2FHA8 | Putative phospholipase | Smlt0142 | 0.007 | 0.43 |
| B2FHB5 | Uncharacterized protein | Smlt0149 | 0.004 | 0.38 |
| B2FHB7 | Glutamine synthetase | glnA | <0.05 | >100 |
| B2FHC6 | Superoxide dismutase [Cu-Zn] | sodC1 | 0.003 | 0.38 |
| B2FHC7 | Superoxide dismutase [Cu-Zn] | sodC2 | 0.032 | 0.49 |
| B2FHD0 | Putative 3-ketoacyl-CoA thiolase | fadI | 0.016 | 0.48 |
| B2FHD6 | Uncharacterized protein | Smlt0170 | 0.007 | 0.41 |
| B2FHE8 | Putative transmembrane protein | Smlt0182 | <0.05 | 2.85 |
| B2FHF0 | Uncharacterized protein | Smlt0184 | 0.006 | 0.39 |
| B2FHF9 | Uncharacterized protein | Smlt0193 | <0.05 | >100 |
| B2FHG4 | Putative electron transfer flavoprotein-ubiquinone oxidoreductase | Smlt0198 | 0.004 | 0.28 |
| B2FHH2 | Putative TonB dependent siderophore receptor | Smlt1426 | 0 | 18.96 |
| B2FHH7 | Putative peptidase | Smlt1431 | 0.012 | 0.36 |
| B2FHI7 | Glutamate--tRNA ligase | gltX | <0.05 | 0.41 |
| B2FHI8 | Putative regulatory protein | fur | <0.05 | >100 |
| B2FHJ2 | Putative outer membrane protein | Smlt1446 | 0.001 | 0.17 |
| B2FHJ5 | Putative phosphodiesterase-nucleotide pyrophosphatase | Smlt1449 | 0.003 | 0.31 |
| B2FHK0 | Uncharacterized protein | Smlt1454 | <0.05 | 0.19 |
| B2FHL6 | Putative ABC transporter | Smlt1471 | 0.019 | 0.38 |
| B2FHL9 | Uncharacterized protein | Smlt1474 | 0.009 | 0.41 |
| B2FHN2 | Uncharacterized protein | Smlt1491 | 0.017 | 0.43 |
| B2FHN5 | Lipid-A-disaccharide synthase | lpxB | 0.013 | 0.48 |
| B2FHQ1 | Putative enterobactin synthetase component A (2,3-dihydro-2,3-dihydroxybenzoate dehydrogenase) | entA | <0.05 | >100 |
| **B2FHQ4** | **Putative hydrolase** | **Smlt2820** | **<0.05** | **>100** |
| B2FHQ5 | Putative siderophore specific 2,3-dihydroxybenzoate-AMP ligase | Smlt2821 | <0.05 | >100 |
| B2FHR5 | Putative transmembrane LINOLEOYL-CoA DESATURASE (DELTA(6)-DESATURASE) | Smlt2831 | <0.05 | 0.42 |
| B2FHR8 | Putative autotransporter protein | Smlt2834 | 0.032 | 0.26 |
| B2FHR9 | Putative TonB-dependent outer membrane protein | Smlt2835 | 0 | 3.57 |
| B2FHT4 | Putative TonB dependent extracellular heme-binding protein | Smlt2850 | <0.05 | >100 |
| B2FHT9 | Putative iron transporter | Smlt2858 | 0 | 4.19 |
| B2FHW5 | Methionine import ATP-binding protein MetN | metN | 0.012 | 0.34 |
| B2FHX5 | Putative CBS domain protein | Smlt4098 | <0.05 | >100 |
| B2FHY7 | ATP synthase epsilon chain | atpC | 0.002 | 0.43 |
| B2FHZ2 | ATP synthase subunit b | atpF | 0.013 | 0.4 |
| B2FHZ7 | Putative dihydrolipoamide dehydrogenase | lpdA | 0.001 | 2.48 |
| B2FHZ8 | Acetyltransferase component of pyruvate dehydrogenase complex | pdhB | 0.012 | 2.71 |
| B2FI00 | Putative outer membrane Omp family protein | Smlt4123 | 0.002 | 0.25 |
| B2FI05 | Conserved hypothetical exported protein | Smlt4128 | 0.006 | 0.45 |
| B2FI10 | Glucans biosynthesis glucosyltransferase H | opgH | <0.05 | >100 |
| B2FI12 | Putative colicin I receptor | cirA | 0 | 75.12 |
| B2FI15 | Putative methyl-accepting chemotaxis protein | Smlt4138 | <0.05 | >100 |
| B2FI22 | Putative extracellular serine protease | Smlt4145 | <0.05 | >100 |
| B2FI29 | Putative NAD(P)H dehydrogenase | Smlt0209 | 0.011 | 0.19 |
| B2FI43 | Putative peptidyl-dipeptidase Dcp (Dipeptidyl carboxypeptidase) | dcp | 0.01 | 0.41 |
| B2FI64 | Ribonucleoside-diphosphate reductase | RRM1 | 0.008 | 3.67 |
| B2FI80 | DntE | Smlt0264 | <0.05 | 0.39 |
| B2FIA0 | Outer membrane protein assembly factor BamA | bamA | 0.01 | 0.48 |
| B2FIA8 | Elongation factor Ts | tsf | 0.015 | 2.07 |
| B2FIC9 | Putative multidrug resistance protein A | emrA | 0.025 | 0.37 |
| B2FIE0 | Putative two-component system, response regulator transcriptional regulatory protein | Smlt1540 | <0.05 | 0.47 |
| B2FII5 | Putative CDP-diacylglycerol pyrophosphatase | Smlt2903 | 0.009 | 0.37 |
| B2FIL8 | Putative TonB dependent receptor protein | Smlt2937 | <0.05 | >100 |
| B2FIM0 | Putative TonB-like protein | Smlt2939 | <0.05 | >100 |
| B2FIQ4 | GTP cyclohydrolase II | Smlt4168 | <0.05 | 0.42 |
| B2FIR5 | Putative LysM family cell wall degradation protein | Smlt4179 | 0.035 | 0.12 |
| B2FIR9 | Putative transmembrane RDD family protein | Smlt4183 | 0.016 | 0.47 |
| B2FIS0 | DNA topoisomerase 1 | topA | <0.05 | >100 |
| B2FIS4 | Putative lipid biosynthesis 3-oxoacyl-[acyl-carrier-protein] reductase | Smlt4188 | <0.05 | 0.46 |
| B2FIS6 | Protease 4 | sppA | 0.002 | 0.38 |
| B2FIS7 | Putative drug resistance transport protein | Smlt4191 | 0.005 | 0.43 |
| B2FIU9 | 60 kDa chaperonin | groL | 0.001 | 2.22 |
| B2FIV0 | 10 kDa chaperonin | groS | 0 | 3.64 |
| B2FJ30 | Putative TonB dependent receptor protein | Smlt0359 | 0.022 | 0.33 |
| B2FJ60 | Putative transmembrane PepSY domain protein | Smlt1566 | <0.05 | >100 |
| B2FJ77 | Polyamine-transporting ATPase | potG | 0.013 | 0.36 |
| B2FJ78 | Putative putrescine transport system permease protein | potH | 0.007 | 0.32 |
| B2FJ81 | Putative transmembrane magnesium/cobalt transport protein | corA | 0.039 | 0.49 |
| B2FJB1 | Putative oar family adhesion protein | Smlt1619 | 0.002 | 0.32 |
| B2FJB8 | Putative PilE protein (Type 4 fimbrial biogenesis protein PilE) | pilE | 0.033 | 0.43 |
| B2FJC0 | UvrABC system protein B | uvrB | <0.05 | 0.43 |
| B2FJC8 | Histidine kinase | Smlt1636 | 0.03 | 0.47 |
| B2FJG0 | Conserved hypothetical exported protein | Smlt2989 | 0.027 | 0.42 |
| B2FJJ3 | Putative hydroxamate-type ferrisiderophore receptor | Smlt3022 | 0 | 22.5 |
| B2FJP1 | Uncharacterized protein | Smlt4245 | 0.017 | 0.44 |
| B2FJQ7 | Putative cell division ATP-binding protein FtsE | ftsE | 0.013 | 0.39 |
| B2FJR1 | Putative transmembrane GGDEF signalling protein | Smlt4266 | 0.021 | 0.48 |
| B2FJR8 | Putative isocitrate dehydrogenase [NADP] | icd | 0.002 | 4.57 |
| B2FJS3 | Putative multidrug efflux system HlyD family transmembrane protein | Smlt4279 | 0.007 | 0.36 |
| B2FJS5 | Putative AcrB/AcrD/AcrF family protein | Smlt4281 | <0.05 | >100 |
| B2FJV0 | Uncharacterized protein | Smlt0387 | 0.002 | 0.39 |
| B2FJV8 | Putative oxidoreductase | Smlt0396 | <0.05 | 0.49 |
| B2FJV9 | Uncharacterized protein | Smlt0397 | 0.02 | 0.48 |
| B2FJW5 | Putative phosphomannomutase | spgM | 0.022 | 0.42 |
| B2FK31 | Putative ABC transporter ATP-binding protein | Smlt1653 | <0.05 | >100 |
| B2FK68 | Putative glutamate synthase | Smlt1693 | 0.003 | 0.22 |
| B2FK78 | Putative pit accessory protein | Smlt1705 | 0.02 | 0.46 |
| B2FK79 | Putative LOW-AFFINITY INORGANIC PHPHATE TRANSPORTER INTEGRAL MEMBRANE PROTEIN PITA | pitA | 0.022 | 0.41 |
| B2FK80 | Putative carboxypeptidase | Smlt1707 | <0.05 | >100 |
| B2FK88 | Enolase | eno | <0.05 | 12.72 |
| B2FK97 | Putative subfamily M23B unassigned peptidase | Smlt1724 | 0.003 | 0.36 |
| B2FKE9 | Putative P-protein [bifunctional includes: chorismate mutase and prephenate dehydratase | pheA | 0.012 | 0.08 |
| B2FKF1 | Conserved hypothetical FHA domain protein | Smlt3099 | 0.021 | 0.49 |
| B2FKG5 | Putative TonB-dependent receptor | Smlt3115 | 0.01 | 0.47 |
| B2FKH5 | Putative transmembrane protein | Smlt3125 | <0.05 | 0.35 |
| B2FKJ7 | 50S ribosomal protein L9 | rplI | 0.003 | 2.68 |
| B2FKJ9 | 30S ribosomal protein S6 | rpsF | 0.003 | 2.62 |
| B2FKK8 | Putative cyclic AMP receptor protein,catabolite gene activator | cap | 0.001 | 2.43 |
| B2FKM2 | Putative K(+)/H(+) antiporter subunit A/B (PH adaptation potassium efflux system protein A/B) | phaAB | <0.05 | 0.35 |
| B2FKR5 | Putative nitrogen regulatory protein P-II | glnB | <0.05 | >100 |
| B2FKV5 | Uncharacterized protein | Smlt0481 | <0.05 | 0.47 |
| B2FKV6 | Putative NADH dehydrogenase/NAD(P)H nitroreductase Smlt0482 | Smlt0482 | <0.05 | 2.24 |
| B2FKW2 | Pyruvate dehydrogenase E1 component | Smlt0490 | 0.01 | 3.25 |
| B2FKX4 | Uncharacterized protein | Smlt0503 | <0.05 | >100 |
| B2FL00 | Putative NHL repeat protein | Smlt0528 | 0.027 | 0.42 |
| B2FL08 | Putative transmembrane anchor protein | Smlt0538 | 0.002 | 0.22 |
| B2FL10 | Uncharacterized protein | Smlt0540 | 0.002 | 0.26 |
| B2FL11 | Putative aminopeptidase | Smlt0541 | <0.05 | 0.23 |
| B2FL46 | Putative cytochrome c family protein | Smlt1757 | <0.05 | >100 |
| B2FL47 | Putative alcohol dehydrogenase cytochrome c subunit | adhB | <0.05 | >100 |
| B2FL51 | Putative ferric siderophore receptor protein | Smlt1762 | <0.05 | >100 |
| B2FL84 | Putative succinate dehydrogenase cytochrome b-556 subunit | sdhC | 0.002 | 0.26 |
| B2FL86 | Putative succinate dehydrogenase flavoprotein subunit | sdhA | 0.006 | 0.2 |
| B2FL87 | Succinate dehydrogenase iron-sulfur subunit | sdhB | 0.002 | 0.23 |
| B2FLA6 | Putative RND family acriflavine resistance protein A | smeG | 0.011 | 0.48 |
| B2FLA7 | Putative multidrug resistance efflux pump | smeH | 0.02 | 0.41 |
| B2FLB8 | Peptidyl-prolyl cis-trans isomerase | Smlt3182 | 0.005 | 0.35 |
| B2FLC5 | Conserved hypothetical exported protein | Smlt3189 | <0.05 | 2.04 |
| B2FLD1 | Putative 2-oxoglutarate dehydrogenase E1 component | sucA | 0.027 | 2.6 |
| B2FLE4 | Putative outer membrane antigen protein | Smlt3210 | 0.036 | 0.49 |
| B2FLE9 | Putative outer membrane antigen lipoprotein | Smlt3215 | 0.015 | 0.45 |
| B2FLF2 | Putative phospholipase | Smlt3218 | 0.005 | 0.41 |
| B2FLG5 | Uncharacterized protein | Smlt3232 | 0.006 | 0.4 |
| B2FLG9 | Putative peptide transport protein | Smlt3236 | 0.029 | 0.48 |
| B2FLH1 | Superoxide dismutase | sodA | 0 | 2.77 |
| B2FLH4 | Putative transmembrane repetitive protein | Smlt3241 | 0.027 | 4.81 |
| B2FLP0 | Putative transmembrane Na+/H+ antiport transporter | Smlt0555 | 0.038 | 0.49 |
| B2FLQ1 | D-amino acid dehydrogenase | dadA | 0.016 | 0.32 |
| B2FLR8 | Putative vitamin B12 receptor protein | Smlt0585 | 0.006 | 0.48 |
| B2FLT1 | Asparagine synthetase | Smlt0598 | <0.05 | >100 |
| B2FLU8 | Transport permease protein | wzm | 0.021 | 0.49 |
| B2FLV4 | Putative transmembrane protein | Smlt0624 | 0.041 | 0.49 |
| B2FLV6 | Putative transmembrane protein | Smlt0626 | 0.01 | 0.34 |
| B2FLW2 | Chaperone protein HtpG | htpG | 0.007 | 4.25 |
| B2FLW5 | Conserved hypothetical exported protein | Smlt1812 | 0.015 | 0.46 |
| B2FLX9 | Putative outer membrane lipoprotein | Smlt1826 | 0.011 | 0.43 |
| B2FLZ4 | Uncharacterized protein | Smlt1841 | <0.05 | 0.42 |
| B2FM92 | Ribonuclease E | rnE | 0.023 | 2.36 |
| B2FMB8 | Putative cytochrome C-type biogenesis protein | dsbE | 0.027 | 0.26 |
| B2FMC4 | Putative ABC transporter ATP-binding protein, cytochrome related | cydD | <0.05 | 0.25 |
| B2FMF2 | Uncharacterized protein | Smlt3312 | 0.015 | 0.46 |
| B2FMI1 | Putative transmembrane protein | Smlt4498 | 0.009 | 0.45 |
| B2FMI9 | Putative TonB dependent receptor protein | Smlt4506 | 0.01 | 0.4 |
| B2FMK1 | Putative ABC transporter ATP-binding protein | Smlt4518 | 0.014 | 0.46 |
| B2FML3 | Uncharacterized protein | Smlt4532 | <0.05 | >100 |
| B2FMN5 | Putative transmembrane UbiA prenyltransferase family protein | Smlt0634 | 0.012 | 0.37 |
| B2FMP6 | Putative electron transfer flavoprotein subunit beta | etfS | 0.001 | 3.18 |
| B2FMP8 | Glucose-1-phosphate thymidylyltransferase | rmlA | <0.05 | >100 |
| B2FMQ1 | Putative transmembrane protein | Smlt0651 | 0.021 | 0.39 |
| B2FMR2 | Putative permease component of ABC transporter protein | Smlt0662 | 0 | 0.28 |
| B2FMR3 | Putative transmembrane protein | Smlt0663 | 0.002 | 2.24 |
| B2FMR8 | Putative transmembrane CDP-diacylglycerol--serine O-phosphatidyltransferase | Smlt0669 | 0.023 | 0.44 |
| B2FMS5 | Putative transmembrane permease protein | Smlt0676 | 0.008 | 0.39 |
| B2FMS6 | Putative transmembrane permease protein | Smlt0677 | 0.026 | 0.45 |
| B2FMX9 | Putative ferric uptake regulation protein | fur | <0.05 | 2.61 |
| B2FN02 | Protein translocase subunit SecD | secD | 0.016 | 0.41 |
| B2FN03 | Protein-export membrane protein SecF | secF | 0.016 | 0.47 |
| B2FN25 | Putative carbon-nitrogen hydrolase | Smlt2038 | <0.05 | >100 |
| B2FN37 | Putative N-acetylmuramoyl-L-alanine amidase | Smlt3330 | 0.006 | 0.29 |
| B2FN47 | Putative TonB dependent receptor | Smlt3340 | 0.003 | 0.27 |
| B2FN55 | Putative acyl-CoA dehydrogenase oxidoreductase protein | Smlt3352 | <0.05 | >100 |
| B2FN59 | Conserved hypothetical repetitive protein | Smlt3358 | 0.006 | 0.34 |
| B2FN90 | Translation initiation factor IF-2 | infB | 0.042 | 2.48 |
| B2FN93 | NADH-quinone oxidoreductase subunit N | nuoN | 0.022 | 0.27 |
| B2FN94 | Putative NADH dehydrogenase I chain M | nuoM | 0.016 | 0.21 |
| B2FN95 | Putative NADH-ubiquinone oxidoreductase I chain L | nuoL | 0.02 | 0.25 |
| B2FNA4 | Putative exported lipoprotein | Smlt4560 | 0.029 | 0.21 |
| B2FNA6 | Putative transmembrane protein | Smlt4562 | 0.021 | 0.48 |
| B2FNC5 | Putative dipeptidyl peptidase | Smlt4581 | 0.004 | 0.42 |
| B2FND7 | Uncharacterized protein | Smlt4593 | 0.002 | 0.3 |
| B2FNF1 | Putative transmembrane efflux pump protein | smmQ | <0.05 | 0.38 |
| B2FNF4 | Putative glycosyl transferase | Smlt4611 | 0.003 | 0.26 |
| B2FNG6 | Acetyl-coenzyme A synthetase | acsA | <0.05 | >100 |
| B2FNK2 | Serine hydroxymethyltransferase | glyA | <0.05 | >100 |
| B2FNL0 | Uncharacterized protein | Smlt0726 | 0.045 | 0.44 |
| B2FNN2 | Putative penicillin-binding protein 3 | ftsI | 0.005 | 0.37 |
| B2FNN3 | UDP-N-acetylmuramoyl-L-alanyl-D-glutamate--2,6-diaminopimelate ligase | murE | <0.05 | 0.27 |
| B2FNN7 | UDP-N-acetylglucosamine--N-acetylmuramyl-(pentapeptide) pyrophosphoryl-undecaprenol N-acetylglucosamine transferase | murG | 0.012 | 0.45 |
| B2FNP5 | 30S ribosomal protein S1 | rpsA | 0.001 | 3.19 |
| B2FNQ8 | Putative transmembrane protein | Smlt2057 | 0.045 | 0.46 |
| B2FNQ9 | Putative transmembrane protein | Smlt2058 | 0.02 | 0.36 |
| B2FNS0 | Inosine-5'-monophosphate dehydrogenase | guaB | 0.005 | 5.15 |
| B2FNX0 | Putative NADH-ubiquinone oxidoreductase, chain J | nuoJ | <0.05 | 0.15 |
| B2FNX1 | NADH-quinone oxidoreductase subunit I | nuoI | 0.004 | 0.23 |
| B2FNX2 | NADH-quinone oxidoreductase subunit H | nuoH | 0.005 | 0.11 |
| B2FNX3 | Putative NADH-ubiquinone oxidoreductase, 75 kDa subunit | nuoG | 0.009 | 0.19 |
| B2FNX4 | Putative NADH dehydrogenase I chain F | nuoF | 0.019 | 0.19 |
| B2FNX5 | Putative respiratory-chain NADH dehydrogenase I, 24 kDa subunit | nuoE | <0.05 | >100 |
| B2FNX6 | NADH-quinone oxidoreductase subunit D | nuoD | 0.003 | 0.32 |
| B2FNX7 | NADH-quinone oxidoreductase subunit C | nuoC | 0.001 | 0.21 |
| B2FNX8 | NADH-quinone oxidoreductase subunit B | nuoB | 0.002 | 0.23 |
| B2FNY0 | Putative general secretory pathway protein-export membrane protein | secG | 0.002 | 0.42 |
| B2FNY5 | Putative lipopolysaccharide core oligosaccharide biosynthesis protein | Smlt3412 | 0.01 | 0.31 |
| B2FNZ9 | Aspartate-semialdehyde dehydrogenase | asd | <0.05 | 0.44 |
| B2FP18 | Putative endopeptidase O | pepO | 0.019 | 0.48 |
| B2FP46 | Putative surface antigen exported protein | Smlt4633 | 0.021 | 0.49 |
| B2FP53 | Sec-independent protein translocase protein TatB | tatB | 0.008 | 0.4 |
| B2FP71 | GTP cyclohydrolase 1 | folE | <0.05 | >100 |
| B2FP87 | Putative intercellular spreading VacJ lipoprotein | Smlt4675 | 0.011 | 0.47 |
| B2FP99 | Uncharacterized protein | Smlt4687 | <0.05 | >100 |
| B2FPA4 | Putative polysaccharide deacetylase family protein | Smlt4692 | 0.012 | 0.41 |
| B2FPA5 | Membrane protein insertase YidC | yidC | 0.01 | 0.46 |
| B2FPC4 | Uncharacterized protein | Smlt0777 | <0.05 | 0.19 |
| B2FPD1 | Putative glycosyl transferase | Smlt0784 | 0.019 | 0.47 |
| B2FPD9 | Probable Na+ dependent nucleoside transporter | Smlt0792 | 0.001 | 0.44 |
| B2FPE2 | Putative exported heme receptor protein | huvA | 0 | 42.38 |
| B2FPE3 | Uncharacterized protein | Smlt0796 | <0.05 | >100 |
| B2FPE4 | Uncharacterized protein | Smlt0797 | <0.05 | >100 |
| B2FPF2 | Putative transmembrane protein | Smlt0805 | 0.006 | 0.39 |
| B2FPF9 | Prolipoprotein diacylglyceryl transferase | lgt | 0.01 | 0.43 |
| B2FPH9 | Glutamine--tRNA ligase | glnS | <0.05 | 0.08 |
| B2FPK0 | Putative repetitive protein with two-component sensor and regulator motifs | Smlt2141 | 0.012 | 0.24 |
| B2FPM6 | Putative calcineurin phosphoesterase | Smlt2170 | <0.05 | 0.48 |
| B2FPN5 | Putative TonB-dependent outer membrane receptor protein | Smlt2179 | 0.006 | 0.23 |
| B2FPP8 | Putative amino-acid transporter transmembrane protein | Smlt2193 | 0.002 | 0.4 |
| B2FPQ0 | Putative D-3-phosphoglycerate dehydrogenase | serA | <0.05 | >100 |
| B2FPR6 | LPS-assembly lipoprotein LptE | lptE | 0.008 | 0.46 |
| B2FPS0 | Putative exported LysM bacterial cell wall related protein | Smlt3488 | 0.004 | 0.28 |
| B2FPS1 | Putative transmembrane GGDEF transcriptional regulatory protein | Smlt3489 | <0.05 | >100 |
| B2FPV6 | Putative thiolase | Smlt3525 | <0.05 | 0.37 |
| B2FPY0 | Uncharacterized protein | Smlt3550 | 0.007 | 0.38 |
| B2FPY7 | Putative alkyl hydroperoxide reductase subunit c | ahpC | 0 | 7.71 |
| B2FPY9 | Proline iminopeptidase | pip | <0.05 | >100 |
| B2FQ15 | Glutamyl-tRNA reductase | hemA | <0.05 | 0.18 |
| B2FQ17 | Outer-membrane lipoprotein LolB | lolB | 0.038 | 0.48 |
| B2FQ28 | Putative TonB dependent receptor protein | Smlt0885 | 0.005 | 0.41 |
| B2FQ31 | Elongation factor Tu | tufB | 0.001 | 3.65 |
| B2FQ33 | Transcription termination/antitermination protein NusG | nusG | 0.036 | 2.41 |
| B2FQ36 | 50S ribosomal protein L10 | rplJ | 0.001 | 2.54 |
| B2FQ38 | DNA-directed RNA polymerase subunit beta | rpoB | 0.013 | 5.3 |
| B2FQ39 | DNA-directed RNA polymerase subunit beta' | rpoC | 0.006 | 7.89 |
| B2FQ42 | Elongation factor G | fusA | 0.003 | 4.86 |
| B2FQ57 | Uncharacterized protein | Smlt2204 | 0.049 | 0.47 |
| B2FQ62 | Putative enoyl-CoA hydratase/isomerase | Smlt2209 | <0.05 | 0.44 |
| B2FQ73 | Transcription elongation factor GreA | greA | <0.05 | >100 |
| B2FQ89 | Oxygen-dependent choline dehydrogenase | betA | 0.007 | 0.35 |
| B2FQD9 | Putative transmembrane protein | Smlt3569 | <0.05 | >100 |
| B2FQE1 | Putative transmembrane sulfatase | Smlt3571 | 0.004 | 0.22 |
| B2FQE3 | Putative angiotensin-converting enzyme like peptidyl dipeptidase protein | Smlt3574 | 0.018 | 0.37 |
| B2FQH7 | Citrate synthase | prpC | <0.05 | 0.38 |
| B2FQJ2 | Putative two-component histidine kinase/response regulator fusion protein | Smlt3625 | 0.04 | 0.41 |
| B2FQK3 | 50S ribosomal protein L15 | rplO | 0.006 | 2.18 |
| B2FQK4 | Protein translocase subunit SecY | secY | 0.015 | 0.4 |
| B2FQK8 | DNA-directed RNA polymerase subunit alpha | rpoA | 0.002 | 3.72 |
| B2FQL7 | Peptidyl-prolyl cis-trans isomerase | ppi | 0.003 | 2.1 |
| B2FQL8 | Malate dehydrogenase | mdh | 0 | 3.96 |
| B2FQM3 | Putative glutathione S-transferase | gst | <0.05 | >100 |
| B2FQN2 | Putative 2-amino-3-ketobutyrate coenzyme A ligase | kbl | <0.05 | 2.22 |
| B2FQN3 | Uncharacterized protein | Smlt0960 | 0.003 | 0.33 |
| B2FQN4 | L-threonine 3-dehydrogenase | tdh | <0.05 | >100 |
| B2FQQ5 | Putative isocitrate/isopropylmalate dehydrogenase | Smlt0982 | 0.001 | 4.75 |
| B2FQR3 | ATP-dependent Clp protease ATP-binding subunit ClpX | clpX | 0.025 | 2.39 |
| B2FQS5 | Putative flagellar basal body-associated protein FliL | fliL | <0.05 | 0.25 |
| B2FQU5 | Putative flagellin | fliC | 0.038 | 2.62 |
| B2FQU6 | Putative flagellin | flaA | 0.034 | 2.68 |
| B2FQY3 | Thioredoxin reductase | trxB | <0.05 | >100 |
| B2FR07 | Uncharacterized protein | Smlt3644 | <0.05 | >100 |
| B2FR21 | Aminomethyltransferase | gcvT | <0.05 | 0.27 |
| B2FR42 | Conserved hypothetical exported protein | Smlt3680 | 0.024 | 0.42 |
| B2FR45 | Putative O-antigen related protein | wbpV | 0.017 | 0.46 |
| B2FR62 | Putative autotransporter | Smlt1001 | <0.05 | 0.04 |
| B2FR74 | Putative DNA polymerase III subunit Tau | dnaX | <0.05 | 0.48 |
| B2FR82 | Putative transmembrane protein | Smlt1022 | 0.001 | 0.2 |
| B2FR90 | Acyl carrier protein | acpP | <0.05 | >100 |
| B2FR91 | 3-oxoacyl-[acyl-carrier-protein] synthase 2 | fabF | <0.05 | >100 |
| B2FRC4 | Putative TonB-dependent receptor | Smlt1067 | 0.024 | 3.11 |
| **B2FRE3** | **Putative esterase** | **Smlt2353** | **<0.05** | **>100** |
| **B2FRE4** | **Putative ATP-binding protein** | **Smlt2354** | **<0.05** | **>100** |
| **B2FRE5** | **Putative binding-protein-dependent transport lipoprotein** | **Smlt2355** | **0** | **19.84** |
| **B2FRE6** | **Putative FecCD-family transmembrane transport protein** | **Smlt2356** | **<0.05** | **>100** |
| **B2FRE7** | **Hemin import ATP-binding protein HmuV** | **hmuV** | **<0.05** | **>100** |
| B2FRM8 | Putative peptidoglycan-associated lipoprotein | Smlt3703 | 0.005 | 0.42 |
| B2FRN1 | Putative TolR-related protein | tolR | 0.013 | 0.42 |
| B2FRN2 | Putative TolQ transport transmembrane protein | tolQ | 0.018 | 0.35 |
| B2FRR0 | Putative lipoprotein | Smlt3739 | 0.001 | 0.39 |
| B2FRU0 | Pyrroline-5-carboxylate reductase | Smlt1087 | <0.05 | >100 |
| B2FRV6 | Putative MgtE/divalent cation transmembrane transporter protein | Smlt1103 | 0.028 | 0.47 |
| B2FRW5 | Putative ABC transporter, ATP-binding protein | Smlt1113 | 0.014 | 0.5 |
| B2FRX9 | Conserved hypothetical exported protein | Smlt1127 | 0.013 | 0.49 |
| B2FRZ2 | Conserved hypothetical exported protein | Smlt1140 | <0.05 | 14.83 |
| B2FRZ3 | Conserved hypothetical exported protein | Smlt1141 | <0.05 | >100 |
| B2FRZ9 | Putative iron transport receptor protein | Smlt1148 | 0 | 2.21 |
| B2FS06 | Putative ABC transporter, ATP-binding protein | Smlt1155 | <0.05 | 2.41 |
| B2FS15 | Putative histone-like protein | Smlt1164 | 0 | 4.96 |
| B2FS22 | Putative copper-transporting p-type ATPase | copF | <0.05 | 0.17 |
| B2FSD2 | Putative transmembrane transporter | Smlt3777 | 0.014 | 0.41 |
| B2FSE0 | Putative transmembrane symporter | Smlt3785 | 0.017 | 0.36 |
| B2FSE3 | Putative HlyD-family secretion protein | smeM | 0.001 | 0.37 |
| B2FSE4 | Putative TonB dependent receptor | Smlt3789 | 0.006 | 2.18 |
| B2FSE9 | Conserved hypothetical exported protein | Smlt3796 | 0.001 | 0.28 |
| B2FSF0 | Fructose-bisphosphate aldolase | alf1 | 0 | 2.71 |
| B2FSF3 | Phosphoglycerate kinase | pgk | <0.05 | 3.06 |
| B2FSF6 | Glyceraldehyde-3-phosphate dehydrogenase | gap | 0.001 | 3.04 |
| B2FSF7 | Putative outer membrane Omp family protein | Smlt3805 | 0.045 | 0.1 |
| B2FSF8 | Putative endonuclease P1 | Smlt3806 | 0.004 | 0.45 |
| B2FSG4 | Uncharacterized protein | Smlt3812 | 0.016 | 0.38 |
| B2FSG6 | Putative transmembrane sodium-dicarboxylate family transporter protein | Smlt3814 | 0.025 | 0.42 |
| B2FSH0 | Putative transmembrane protein | Smlt3818 | <0.05 | 0.46 |
| B2FSH4 | Putative pilP protein (Type 4 fimbrial biogenesis protein) | pilP | <0.05 | 0.32 |
| B2FSH6 | Putative PilN protein (Type 4 fimbrial biogenesis protein) | pilN | <0.05 | 0.35 |
| B2FSH7 | Putative PilM protein (Type 4 fimbrial biogenesis protein) | pilM | 0.038 | 0.21 |
| B2FSJ9 | Putative TonB dependent receptor protein | Smlt1175 | <0.05 | >100 |
| B2FSN4 | Putative transmembrane phosphoesterase | Smlt1210 | 0.017 | 0.31 |
| B2FSQ7 | Putative TonB-dependent receptor for Fe(III)-coprogen, Fe(III)-ferrioxamine B and Fe(III)-rhodotrulic acid | Smlt1233 | 0 | 41.01 |
| B2FT31 | Putative ACR family protein | Smlt3869 | 0.014 | 0.44 |
| B2FT45 | Putative transmembrane DedA family protein | Smlt3883 | 0.03 | 0.5 |
| B2FT59 | Putative extracellular heme-binding protein | Smlt3898 | 0 | 8.18 |
| B2FT63 | Inorganic pyrophosphatase | ppa | <0.05 | 2.3 |
| B2FT66 | Putative TonB dependent receptor | Smlt3905 | 0.014 | 0.44 |
| B2FT80 | DNA polymerase III subunit beta | dnaN | <0.05 | >100 |
| B2FT86 | Conserved hypothetical TPR repeat family protein | Smlt0008 | 0 | 4.78 |
| **B2FT87** | **Putative proline-rich TonB dependent receptor protein** | **Smlt0009** | **0.006** | **0.34** |
| B2FTA3 | Putative exported peptidase | Smlt1246 | 0.033 | 2.06 |
| B2FTA8 | Probable septum site-determining protein MinC | minC | <0.05 | 2.27 |
| B2FTD0 | UvrABC system protein A | uvrA | <0.05 | 0.45 |
| B2FTJ0 | Isoleucine--tRNA ligase | ileS | <0.05 | 2.17 |
| B2FTK5 | Putative ferric siderophore receptor | Smlt2650 | <0.05 | 5.09 |
| B2FTN0 | Putative transmembrane protein | Smlt2678 | <0.05 | 0.32 |
| B2FTR3 | Conserved hypothetical exported protein | Smlt2712 | <0.05 | >100 |
| B2FTR4 | Conserved hypothetical exported protein | Smlt2713 | <0.05 | >100 |
| B2FTR5 | Putative TonB dependent protein, possible siderophore receptor | Smlt2714 | <0.05 | >100 |
| B2FTS4 | Uncharacterized protein | pcm | 0.007 | 0.38 |
| B2FTS5 | Protein CyaE | tolC | 0.001 | 0.25 |
| B2FTS8 | Putative lipid A biosynthesis lauroyl acyltransferase | htrB | 0.017 | 0.44 |
| B2FTT7 | Putative porin P (Outer membrane protein d1) | oprP | 0.007 | 0.45 |
| B2FTV4 | Putative glutaredoxin | Smlt3961 | <0.05 | 2.73 |
| B2FTV7 | Conserved hypothetical exported protein | Smlt3964 | 0.024 | 0.38 |
| B2FTY9 | Putative endonuclease/exonuclease/phosphatase family protein | Smlt3995 | <0.05 | 0.36 |
| B2FU12 | Putative iron transporter protein | Smlt0049 | 0.049 | 5.51 |
| B2FU40 | Putative patatin-like phospholipase | Smlt0080 | 0.005 | 0.42 |
| B2FU42 | Putative TonB dependent receptor protein | Smlt0083 | 0.001 | 0.24 |
| B2FU43 | Acid phosphatase | Smlt0084 | 0 | 0.2 |
| B2FU51 | Thymidine kinase | tdk | <0.05 | >100 |
| B2FU87 | Putative transmembrane protein | Smlt1346 | 0.012 | 0.32 |
| B2FU91 | Putative outer membrane autotransporter | Smlt1350 | <0.05 | >100 |
| B2FUA1 | Putative quinol oxidase subunit 1 | qoxB | 0.011 | 0.32 |
| B2FUA6 | Putative chaperone protein HtpG (Heat shock protein HtpG) | htpG | <0.05 | 0.36 |
| B2FUD9 | Putative HlyD family secretion protein | Smlt1406 | 0.011 | 0.43 |
| B2FUF0 | Putative nucleotide sugar transaminase | Smlt1417 | <0.05 | >100 |
| B2FUL1 | Putative respiratory nitrate reductase subunit | narH | 0.018 | 0.28 |
| B2FUN9 | Putative TonB-dependent receptor for Fe(III)-coprogen, Fe(III)-ferrioxamine B and Fe(III)-rhodotrulic acid | fhuE | 0 | 13.58 |
| B2FUR1 | Putative TonB-dependent outer membrane receptor protein | Smlt4026 | 0.001 | 0.13 |
| B2FUT3 | Putative exported rare lipoprotein A | rlpA | 0.001 | 0.27 |
| B2FUT4 | Putative murein hydrolase | mltB | 0.011 | 0.46 |
| B2FUT9 | Putative rod shape-determining protein | mreC | 0.009 | 0.44 |
| B2FUV1 | Putative multidrug resistance outer membrane protein | smeF | 0.002 | 0.39 |
| B2FUV2 | Putative acriflavin resistance protein B | smeE | 0.013 | 0.31 |
| B2FUV3 | Putative acriflavin resistance protein A | smeD | 0.002 | 0.32 |
